# NeuralFieldManifold: Uncovering the Manifold Geometry of Local Field Potentials

**DOI:** 10.64898/2026.10.28.762631

**Authors:** Kasra Fallah, Haoyu Novak Chen, Rudramani Singha, Eunji Kong, Gergely Turi, Attila Losonczy, Erfan Zabeh

## Abstract

Manifold geometry has become a powerful framework for understanding how the brain organizes high-dimensional neural activity into compact, interpretable structure. To date, however, most theoretical and methodological advances in neural manifold analysis have focused on spiking activity, leaving unresolved a fundamental question: What are the geometric principles governing the dynamics of local field potentials (LFPs)—population level signals reflecting integrated sub-threshold and suprathreshold neural activity that are broadly integral to clinical and brain–computer interface settings. Here, we establish, analytically and empirically, that the lag-embedded dynamics of LFP signals lie on a low-dimensional K-torus, where K directly relates to sustained oscillatory modes present in the signal. Modeling LFPs as bounded autoregressive processes, we show that each oscillatory component contributes an independent circular degree of freedom, while aperiodic 1/f structure and noise contribute only geometric thickness around the manifold without altering its topology. To apply this theory to real nonstationary recordings, we introduce DeepLagField, a physics-informed network that jointly estimates time-varying AR structure and effective model order, with formal guarantees that local toroidal geometry is preserved despite non-stationary nature of neuronal processes. Persistent homology analysis on real recordings reveals topological signatures consistent with the predicted toroidal Betti numbers in primate V1 LFP, rodent hippocampal LFP, and mouse cortical EEG, with varying degrees of agreement across recordings and temporal windows. Crucially, decoding analyses across three distinct neural datasets demonstrate that these geometric structures are not merely descriptive but capture behaviorally relevant information rivaling conventional spectral features. Together, these findings establish a theoretically grounded and empirically validated geometric framework for neural field analysis, extending the manifold perspective beyond spiking activity and offering a complementary language to spectral analysis for understanding and translating population dynamics across neural modalities.

## 1 Introduction

Advances in large-scale neural recording technologies have fundamentally reshaped systems neuroscience, producing high-dimensional datasets that challenge traditional single-neuron analyses. A widely adopted framework for interpreting these recordings is the *manifold hypothesis*: despite the large number of recorded neurons, population activity often occupies a lower-dimensional structure within the high-dimensional observation space Cunningham and Yu [2014]. From this perspective, neural computation can be studied as coordinated population dynamics evolving along neural mani-folds, rather than solely through the activity of individual neurons Vyas et al. [2020]. Theoretical and methodological advances have provided tools for estimating and interpreting these structures in population recordings Gao et al. [2017], Yu et al. [2009], establishing manifold analysis as a central approach to studying neural population dynamics.

Yet this geometric perspective has been developed primarily for spiking activity Levy et al. [2023], Chung et al. [2016], Cohen et al. [2020], Chung and Abbott [2021], leaving a complementary class of population signals comparatively unexplored: local field potentials (LFPs). LFPs arise largely from spatially integrated synaptic and other transmembrane currents, reflecting aspects of shared input and subthreshold population dynamics that are not directly captured by spike counts Buzsáki et al. [2012], Einevoll et al. [2019], Jalaldoust and Zabeh [2025]. Their oscillatory activity has been associated with mechanisms of attention, working memory, and cognitive control Miller et al. [2018], Siegel et al. [2012], Zabeh et al. [2023]. These signals therefore offer an important opportunity to extend geometric analyses beyond spiking activity. Recent advances in field potentials and their spatial characteristics (traveling waves) demonstrate that neural field activity can exhibit coordinated spatiotemporal organization Muller et al. [2018], Jung et al. [2025], Das et al. [2025]; whether such organization gives rise to specific intrinsic dimensions and topologies in delay-coordinate representations, however, remains unresolved.

Here, we develop an analytical framework linking neural field dynamics to the geometry of their lag embeddings. Motivated by the bounded amplitudes commonly observed in finite LFP recordings and by spectra comprising narrowband oscillations superimposed on an aperiodic 1*/f*^*α*^ background Donoghue et al. [2020], Buzsáki and Draguhn [2004], we adopt a tractable autoregressive (AR) description of temporal dynamics Stoica et al. [2005], Kaminski and Blinowska [1991]. In an idealized, locally stationary oscillatory model, each sustained, nonresonant complex-conjugate mode contributes an independent circular phase coordinate, so *K* such modes generate a toroidal phase skeleton T^*K*^ = (*S*^1^)^*K*^ in a sufficiently informative delay embedding. This yields an explicit connection between AR pole structure, oscillatory phase variables, and manifold topology. Damping, stochastic fluctuations, and aperiodic activity can thicken or distort the embedded trajectories, making the predicted topology more or less detectable rather than guaranteeing an exact torus in every observed window. To address finite and nonstationary conditions of real neuronal recording, we extend the analysis to time-varying AR dynamics and derive a local approximation bound. A Lipschitz condition on the coefficient path bounds the mismatch between a TVAR window and its frozen AR surrogate by 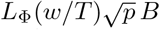, where *L*_Φ_ is the coefficient drift rate, *w* is the window length, *p* is the AR order, and *B* is a local amplitude bound. This condition motivates *DeepLagField*, a physics-informed TVAR estimator whose smoothness and energy regularizers encourage slowly varying coefficients and bounded local signal energy. Detailed comparisons to delay-embedding, topological-neural-manifold, and nonstationary dynamics literature are provided in Appendix A.

We evaluate these predictions in synthetic signals with known dynamics and in neural recordings from primate visual cortex LFP, rodent hippocampal LFP, and mouse cortical EEG. Direct topological investigation using persistent homology analysis on these recordings revelaved topological signatures consistent with the predicted toroidal structures in our theoretical formulation. Beyond geometric characterization, decoding analyses across three datasets show that geometry-derived torus parameters capture behaviorally relevant information with performance comparable to conventional spectral features.

Together, our work establishes a theoretically grounded and empirically tested framework for neural field geometry, extends manifold analysis beyond spiking activity and offering a complementary perspective to spectral analysis for characterizing and decoding population dynamics across neural modalities. Our key contributions are summarized below:

1. **Analytical investigation of manifold geometry for LFP-like signals** We derive an explicit connection between AR pole structure and delay geometry: under ideal noiseless, locally stationary, nonresonant conditions, each sustained conjugate pole pair contributes one circular phase factor, yielding a T^*K*^ toroidal skeleton.
2. **Analytical extensions under nonstationarity: limits and computational motivation** We extend the stationary AR picture to slowly varying TVAR dynamics via a frozen-window approximation bound, giving concrete conditions under which local toroidal structure should remain detectable.
3. **PINN-based solution to joint nonstationary dynamics and order inference** Solving Challenging Order–Dynamics Inference with PINNs We introduce DeepLagField, a physics-informed TVAR estimator that jointly estimates time-varying AR coefficients and effective order while encouraging the smoothness and bounded-energy conditions required for stable geometric recovery.
4. **Validation of topological predictions on real neural recordings** We evaluate the predicted toroidal signatures using persistent homology across primate V1 LFP, rodent hippocampal LFP, and mouse cortical EEG, and compare real embeddings against phase-shuffled surrogates.
5. **Demonstration of behavioral relevance of LFP manifold during tasks** We demonstrate a downstream application of the proposed framework by showing that fitted torus parameters provide reliable, comparable and relevant behavior states on multiple dataset across sessions and animals.

## 2 Analytical Characterization of LFP Manifold Geometry

Local field potentials exhibit bounded amplitude, strong temporal autocorrelations Einevoll et al. [2013], Buzsáki et al. [2012], Lega et al. [2012], Miller et al. [2009], high spatial coherence across electrodes Jung et al. [2025], Muller et al. [2018], Das et al. [2025], and prominent aperiodic spectral feature, often accompanied by oscillatory peaks. Donoghue et al. [2020], Buzsáki and Draguhn [2004]. We show that these characteristics imply a *K*-torus T^*K*^ geometry under delay embedding, where *K* equals the number of distinct sustained oscillatory modes.

### 2.1 Autoregressive model of LFP dynamics

We model a single LFP channel as a stationary autoregressive process of order *p*,

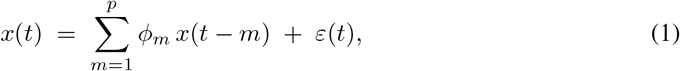

where 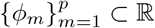 are the AR coefficients and *ε*(*t*) is bounded, zero-mean innovation noise. The multichannel formulation reduces by Lemma B.1 (Appendix B) to single-channel analysis, with each channel contributing independently to the product manifold. The geometry of delay-embedded trajectories is governed entirely by the deterministic component; noise thickens the manifold without altering its topology.

### 2.2 Oscillatory poles, phase variables, and the *K*-torus

The long-run behavior of (1) is determined by the roots of its characteristic polynomial 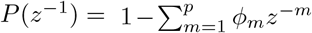 Real coefficients force conjugate pairing. *Unit-circle pairs z* = *e*^*±iω*^, *ω* ∈ (0, *π*), generate sustained oscillations and each introduce one independent phase variable *θ*(*t*) = *ωt* + *φ*_0_ (mod 2*π*) evolving on *S*^1^. *Strictly stable roots* |*z*| *<* 1 contribute only aperiodic background and geometric thickness, without introducing new phase variables (Lemma B.3, Appendix B). Thus *K* sustained modes yield *K* independent circular degrees of freedom.

#### Proposition 2.1

(K-torus embedding; informal). *Let a bounded LFP signal, restricted to a local temporal window, contain exactly K distinct persistent oscillatory components with pairwise distinct frequencies modulo π. In the noiseless marginally-stable limit, a delay embedding using two lags per oscillatory mode lies on a smooth K-dimensional manifold ℳ* ≅ T^*K*^ = (*S*^1^)^*K*^. *Damping, aperiodic structure, and noise appear as geometric thickness around the torus without adding circular factors*.

The formal proof (Proposition B.5, Appendix B) constructs the smooth embedding *F* : T^*K*^ →ℝ^2*K*^ via the realified Vandermonde map and verifies immersion and injectivity via the inverse function theorem.

#### Betti-number corollary

The homology of T^*K*^ satisfies 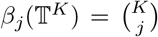 for *j* = 0, …, *K*. In particular *β* = *K* (one circular loop per oscillatory mode) and 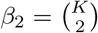. For *K* = 2 — the two-mode case in primate V1 — the predicted signature (*β*_0_, *β*_1_, *β*_2_) = (1, 2, 1) is exactly what persistent homology recovers empirically (Section 3, Figure 5). For *K* = 1 — single-theta rodent hippocampus — (*β*_0_, *β*_1_) = (1, 1) is confirmed likewise. This gives a falsifiable, quantitative prediction directly linking spectral peak count to topological invariants, captured by the correspondence:

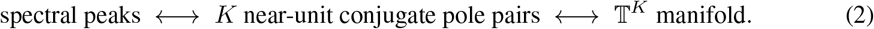

Aperiodic 1*/f*^*α*^ structure and noise affect geometric conditioning — thickening the manifold and introducing anisotropy when oscillatory amplitudes are imbalanced — but do not alter *K* (Appendix J). Linear spectral equalization can improve empirical recoverability without changing topology.

#### Bridge to reconstruction

Realizing this prediction from real recordings poses two challenges: LFPs are nonstationary, violating the fixed-coefficient assumption of (1); and accurate manifold recovery requires correct joint identification of the AR coefficient path *and* the effective order *p*, since low prediction error does not guarantee the correct number of circular factors when *p* is misidentified. Section 3 addresses both through a time-varying AR formulation with a formal local-stationarity guarantee and physics-informed regularization that directly enforces the geometric conditions identified here.

## 3 Reconstruction and Validation

Section 2 establishes that the lag-embedded trajectory of an LFP signal with *K* sustained oscillatory modes lies on T^*K*^. Realising this prediction from experimental recordings requires addressing two challenges absent from the stationary theory: (i) real LFPs are nonstationary, with oscillatory content and amplitudes drifting on behaviorally relevant timescales; and (ii) accurate manifold recovery requires simultaneous correct identification of both the AR coefficient path and the effective model order *p* — since a model achieving low prediction error can nonetheless recover the wrong number of circular factors if *p* is misidentified. We address both challenges in turn.

### 3.1 Local stationarity bridge

We operate in a *locally stationary* regime, modelling the LFP as a time-varying autoregressive (TVAR) process,

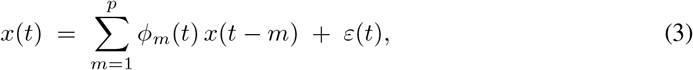

whose coefficient path Φ : [0, 1] → ℝ^*p*^ is Lipschitz with constant *L*_Φ_. The discrepancy between the TVAR process and its frozen AR surrogate at reference time *t*_0_, within a window of length *w*, is bounded by 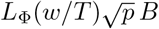, where *B* is the local signal amplitude bound (full derivation in Appendix C). Whenever

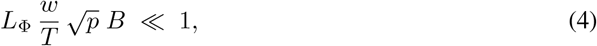

the frozen surrogate is a valid local approximation. Since polynomial roots vary continuously with their coefficients Marden [1966], small coefficient discrepancy implies small pole perturbation — and hence, by Proposition 2.1, the lag-embedded trajectory tracks a T^*K*^ determined by the instantaneous oscillatory modes up to a bounded geometric error. We verify Condition (4) empirically across all three biological datasets by estimating *L*_Φ_ from finite differences of OLS coefficient fits across adjacent windows (Table B.1, Appendix C); the condition holds comfortably in all cases, confirming local stationarity as a valid operating assumption rather than a purely theoretical idealization.

### 3.2 DeepLagField

We introduce *DeepLagField*, a physics-informed network that jointly estimates the time-varying coefficient path 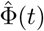 and the effective model order 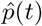 from a TVAR signal. The architecture (Figure 2) consists of two coupled modules.

**Figure 1:**
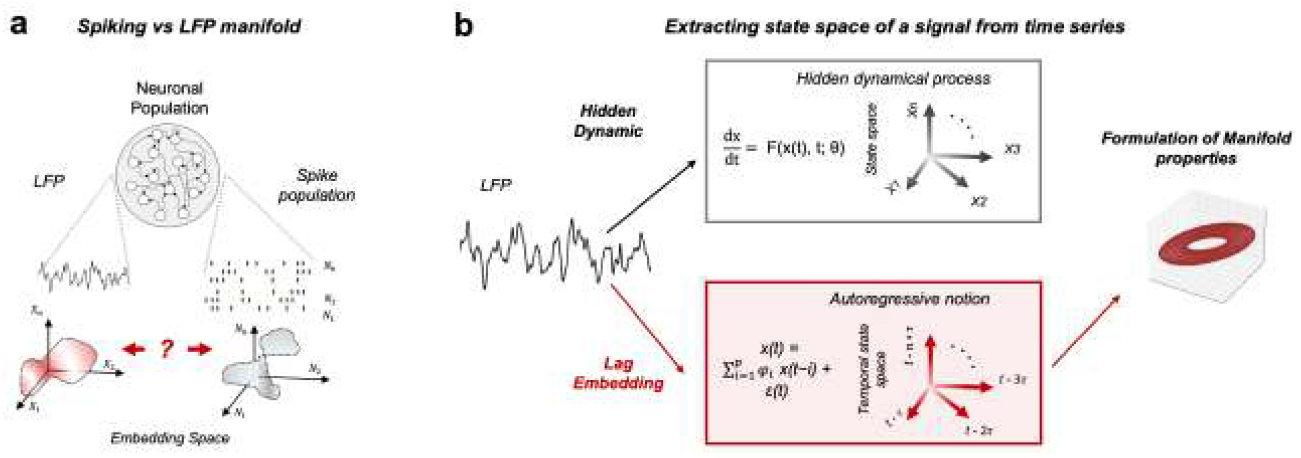
Conceptual framework for deriving LFP manifold geometry. **(a)** While spike-based neural manifolds are well studied, the geometry of aggregate LFP signals remains less understood. **(b)** We formulate LFP time series using autoregressive processes which directly converts these dynamics into a temporal state space.

**Figure 2:**
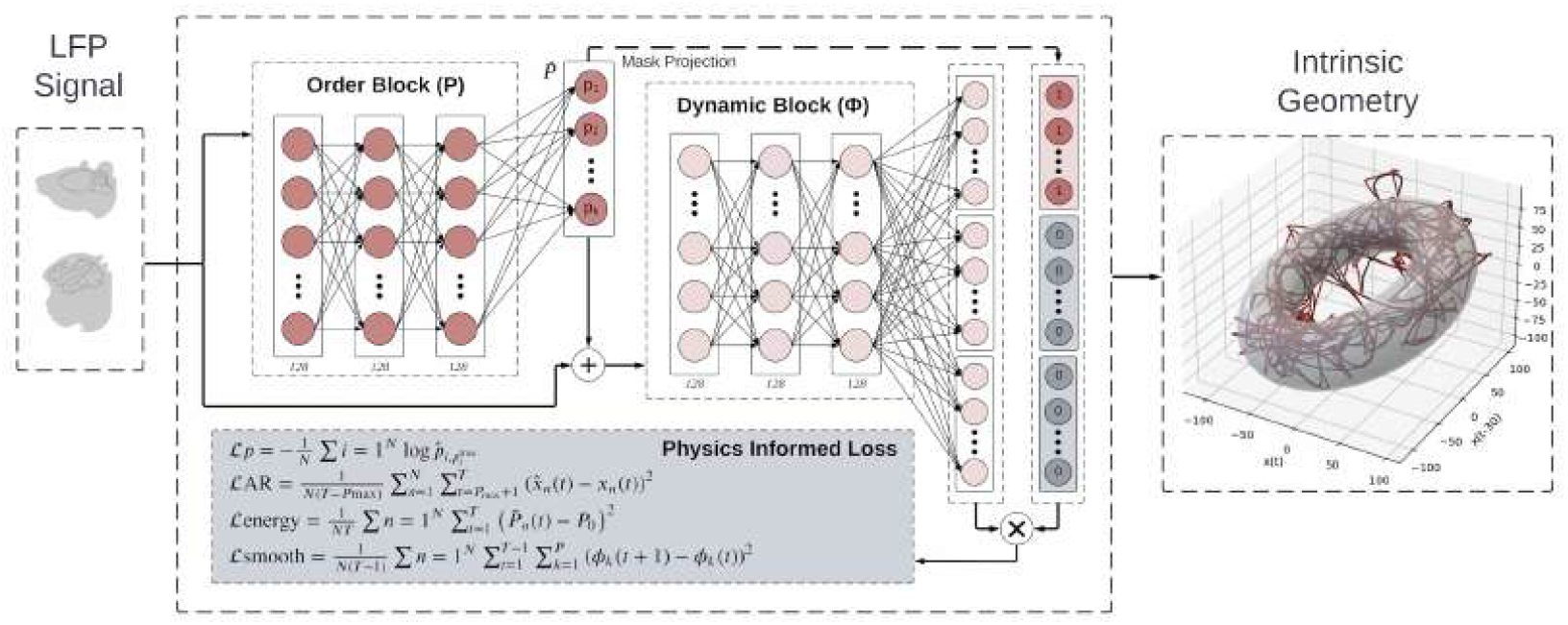
DeepLagField Architecture. The order block produces a sparse mask over candidate lags; the dynamic block, conditioned on this mask, outputs time-varying AR coefficients 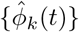. Physics-informed losses enforce prediction fidelity, smoothness of 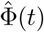, and bounded signal energy.

#### Order block

A lightweight feedforward network maps a short temporal context window to a probability vector over candidate orders {2, …, *p*_max_}, from which a sparse binary mask is obtained via Gumbel-Softmax relaxation with a straight-through estimator Fraser and Swinney [1986]. The resulting effective order 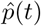 is sparse and time-varying, adapting to the instantaneous oscillatory content of the signal.

#### Dynamic block

Conditioned on the order mask, a second feedforward module outputs time-varying coefficients 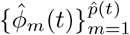. The specific backbone parameterisation is not essential, provided it preserves the lag-based structure required by the theoretical analysis.

#### Physics-informed regularization

The training objective is ℒ = ℒ_pred_ +*λ*_*S*_ ℒ_smooth_ +*λ*_*E*_ ℒ_energy_, where ℒ_pred_ is the one-step prediction MSE, ℒ_smooth_ penalises temporal variation in 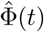 (enforcing small *L*_Φ_, satisfying the numerator of Condition (4)), and ℒ_energy_ penalises rapid fluctuations in local signal power (controlling *B*). Together these regularizers directly operationalize the local-stationarity condition throughout training, rather than imposing it post-hoc. The embedding delay *τ* is selected using the first minimum of average mutual information Fraser and Swinney [1986]. Full training details and hyperparameter specifications are given in Appendix I.

### 3.3 Synthetic validation

We evaluate DeepLagField on synthetic stationary AR and nonstationary TVAR LFP-like (AR(3)) signals as well as on benchmarks with known ground-truth coefficients and order, spanning eight coefficient function classes (sinusoidal, Fourier series, quasi-periodic, polynomial drift, logistic transition, multi-sigmoid, Gaussian bumps, smooth random; details in Appendix G). DeepLagField reconstruction on LFP-like oscillations (Figure 3) shows how predictive accuracy relates to geometric recovery. In both stationary and time-varying AR(3) examples, DeepLagField reconstructs the signal, largely recovers the underlying coefficients, and preserves the lag-embedded manifold structure. OLS achieves comparable prediction accuracy but produces more distorted embeddings, showing why prediction error alone does not establish recovery of geometry.We next examine whether accurate prediction is accompanied by coefficient and order recovery across the broader benchmark suite.Table 1 reports performance averaged across orders *p* ∈ {2, …, 6} and seeds. While the Transformer baseline achieves the lowest prediction MSE (0.013), it fails entirely on coefficient and order recovery — confirming the central claim that prediction accuracy is not a proxy for geometric fidelity. OLS achieves competitive coefficient error but provides no order estimate. DeepLagField achieves the lowest TVAR coefficient error (0.004) and best order stability (0.699) while maintaining competitive prediction performance, demonstrating that the physics-informed constraints successfully decouple reconstruction quality from prediction quality.

**Table 1:**
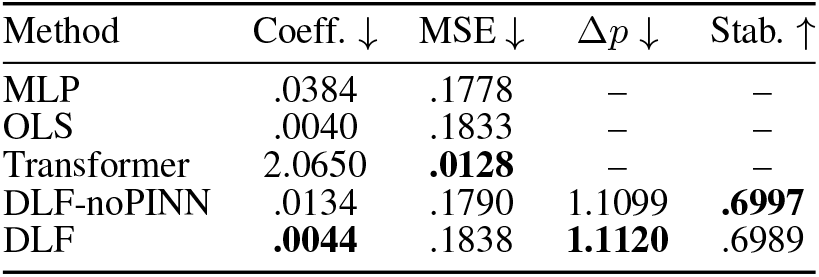
Reconstruction performance on AR and TVAR benchmarks. Mean over orders and seeds. N/A: method does not produce an order

**Figure 3:**
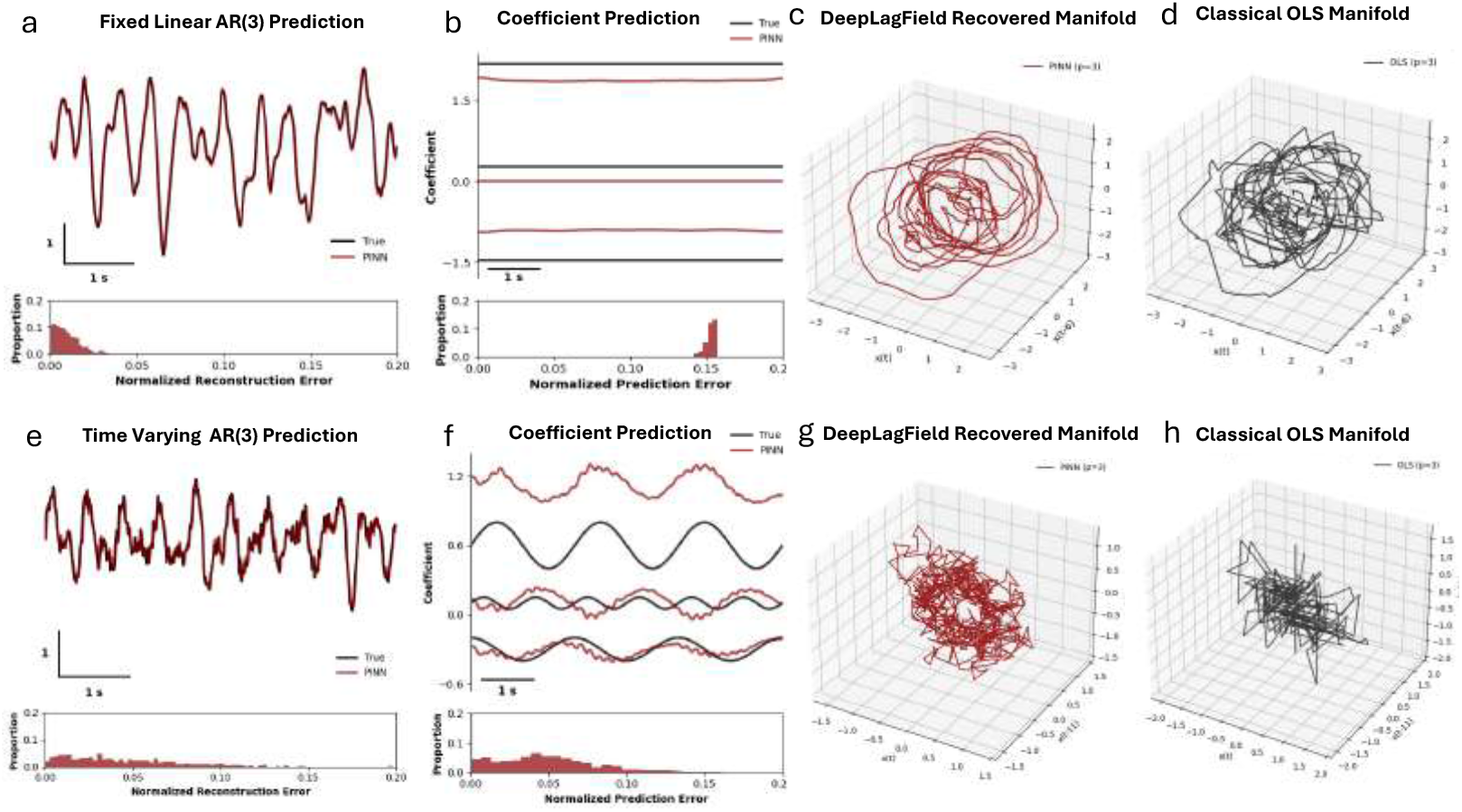
PINN-based reconstruction on synthetic LFP-like signals. Fixed and time-varying AR(3) signals are shown alongside coefficient estimates and manifold embeddings. DeepLagField preserves toroidal structure where OLS collapses geometry despite comparable prediction error .

#### Ablation

Table 2 isolates the contribution of each architectural constraint. Removing the AR structure collapses both coefficient and prediction performance. Removing temporal smoothness sub-stantially degrades coefficient recovery (0.004 →0.016) despite negligible change in prediction error (0.184→ 0.179), confirming that the smoothness regularizer stabilises nonstationary identification independently of predictive accuracy. Most critically, removing the order-consistency constraint leaves prediction error unchanged yet causes order-recovery accuracy to collapse (Δ*p/p* : 0.301 →0.530), demonstrating that correct identification of the number of oscillatory modes — and hence of the torus dimension *K* — requires explicit geometric constraints beyond prediction loss.

**Table 2:**
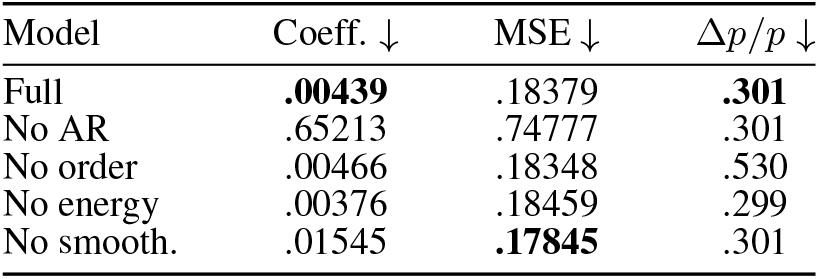
Ablation on TVAR benchmarks. Prediction error alone is insufficient for stable order recovery. Lower ↓ is better.

| Model | Coeff. ↓ | MSE ↓ | $\Delta p/p$ ↓ |
|---|---|---|---|
| Full | **.00439** | .18379 | **.301** |
| No AR | .65213 | .74777 | .301 |
| No order | .00466 | .18348 | .530 |
| No energy | .00376 | .18459 | .299 |
| No smooth. | .01545 | **.17845** | .301 |

### 3.4 Neural data: manifold recovery

We evaluate DeepLagField on three independent electrophysiological datasets: (i) primate V1 LFP (1024-channel Utah array, rhesus macaque, resting state Chen et al. [2022]); (ii) rodent hippocampal LFP (four-channel silicon probe, chronically implanted Grosmark et al. [2021]); and (iii) mouse cortical EEG (chronic skull-screw recordings during sleep Turi et al. [2025]). All analyses use a single pipeline with no dataset-specific tuning; preprocessing details are in Appendix G.

#### Spectral predictions and recovered embeddings

In primate V1, two dominant narrowband peaks in the alpha/beta range predict *K* = 2; in rodent hippocampus, a single dominant theta peak predicts *K* = 1; in mouse cortex, broadband oscillatory activity predicts *K*∈ {1, 2} depending on the recording window. In each case, order selection concentrates at *p*^*\**^ = 2*K* (the minimal embedding dimension from Lemma A.4), and the recovered lag embedding exhibits the predicted toroidal structure. Figure 4 shows the full pipeline for the primate V1 dataset. Cross-species and cross-modality embeddings, including the mouse hippocampal one-torus and the mouse cortex EEG two-torus, are shown in Appendix E (Figures 12, 13). Critically, the same pipeline generalizes across species and brain regions without retuning, indicating that the torus geometry is a modality-invariant property of oscillatory LFP dynamics rather than an artifact of dataset-specific preprocessing.

**Figure 4:**
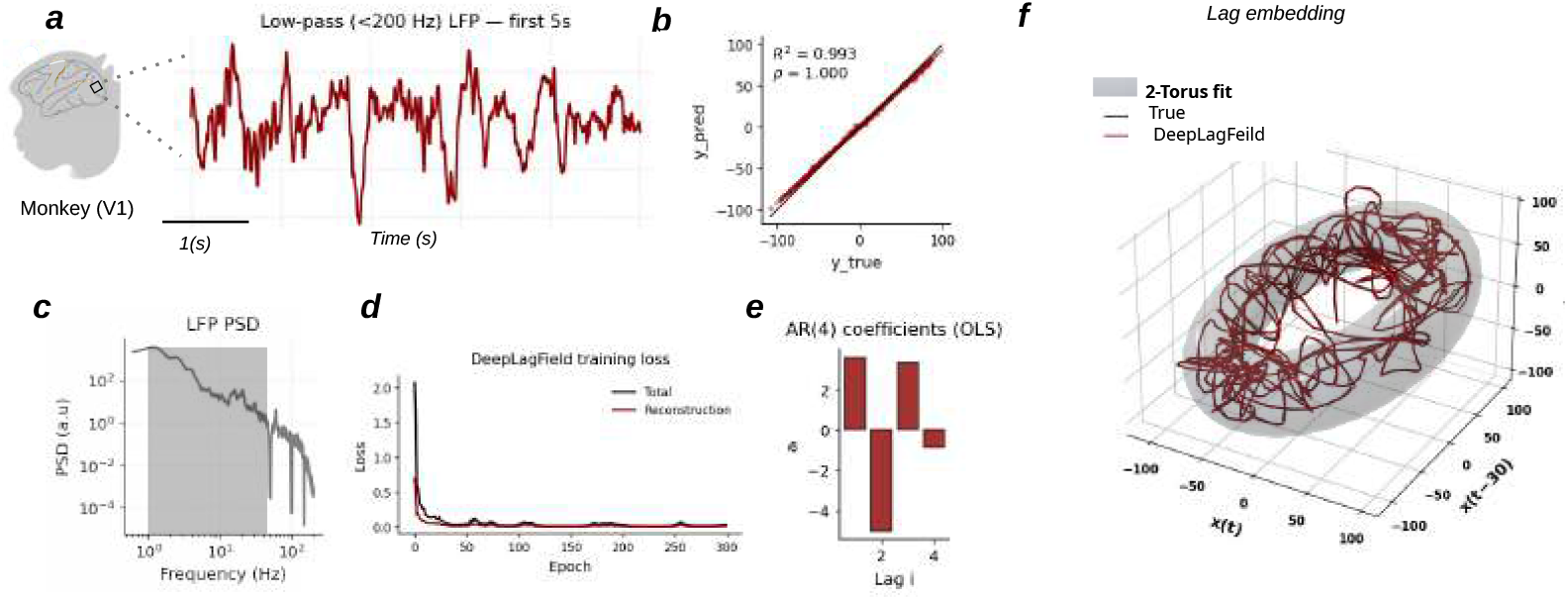
Manifold reconstruction of monkey V1 LFP. **(a)** Example LFP segment from monkey primary visual cortex. **(b)** Held-out prediction performance of DeepLagField, comparing predicted and true samples. **(c)** Power spectral density showing a broadband background with two dominant oscillatory components, predicting *K* = 2. **(d)** Held-out MSE as a function of AR order, selecting *p*^*⋆*^ = 4, consistent with two conjugate pole pairs. **(e)** Estimated time-varying AR coefficients recovered by DeepLagField Bar plots represent coefficient’s mean. **(f)** Delay embedding of the reconstructed dynamics overlaid with a fitted 2-torus, consistent with the predicted toroidal skeleton.

**Figure 5:**
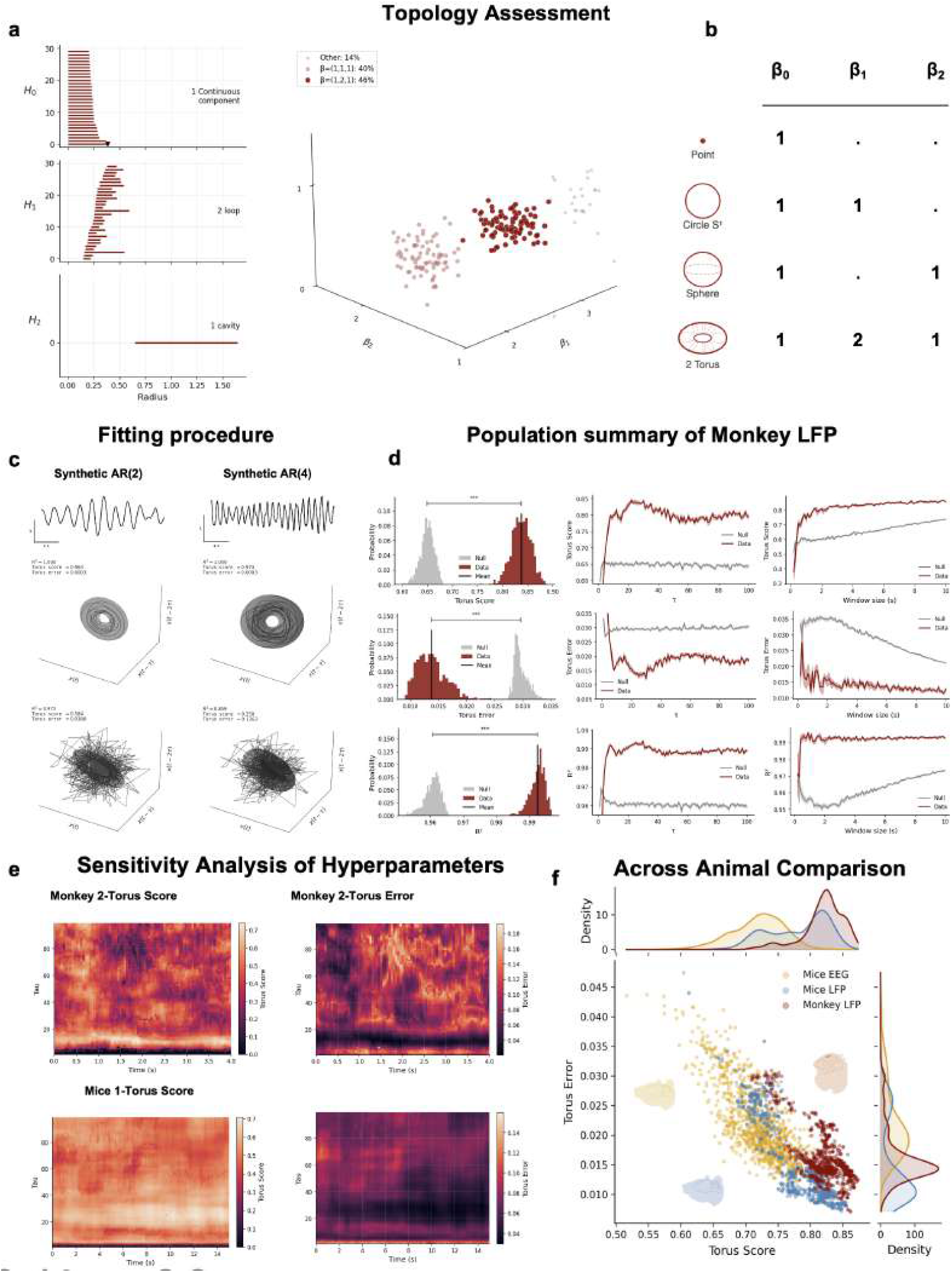
Topological quantification and cross-dataset validation. **(a)** Persistent homology of monkey V1 lag embeddings recovers the predicted two-torus signature (*β*_0_, *β*_1_, *β*_2_) = (1, 2, 1) in the dominant fraction of windows, with partial torus signatures appearing in noisier or less resolved windows. **(b)** Reference Betti numbers for canonical spaces, showing that (1, 2, 1) uniquely matches T^2^ among the displayed geometries. **(c)** Synthetic AR(2) and AR(4) calibration confirms that the pipeline distinguishes T^1^ from T^2^ and does not create toroidal structure from spurious fits. **(d)** Population-level monkey V1 validation: real data show higher torus score and *R*^2^, and lower torus error, than phase-shuffled surrogates (*p <* 0.001). **(e)** Sensitivity analysis shows stable torus scores and errors across embedding delays *τ* and window durations for monkey and mouse recordings. **(f)** Across mouse EEG, mouse LFP, and monkey LFP, stronger toroidal structure consistently corresponds to lower reconstruction error.

### 3.5 Topological validation

We compute persistent homology on lag embeddings of the preprocessed recordings and derive Betti numbers (*β*_0_, *β*_1_, *β*_2_) directly from the point cloud. For primate V1, we sample 200 four-second windows (retaining 300 sample) from the three coordinate lag embedding. After centering and noramlization, Ripser connects nearby points over increasing distance scales and records when loops (*H*_1_) and two-dimensional cavities (*H*_2_) appear and disappear. We classify each window by counting features whose persistence exceeds 0.30. The expected two-torus signature (*β*_0_, *β*_1_, *β*_2_) = (1, 2, 1) appears in 46.5% of windows, (1, 1, 1) in 39.5% of windows, and other signatures in 14%. In the (1, 1, 1) windows, one *H*_1_ class and one *H*_2_ class pass the threshold; the second expected *H*_1_ class does not. We interpret this as a partially resolved toroidal signature, rather than evidence that the manifold collapsed to a circle, which would have signature (1, 1, 0). We compared these window-level Betti classifications with four surrogate controls and one-mode ablation (Table 3) to assess whether the observed split in Betti number distribution is distinctive of the real recording. The (1, 2, 1) signature occurs consistently less often in control windows (23.0–39.0%) than in real signal (46.5%). This supports enrichment at population level while showing that recovery varies across windows. For rodent hippocampa LFP, (*β*_0_, *β*_1_) = (1, 1) is the dominant configuration, consistent with T^1^. Against synthetic true- and spurious-torus controls (Appendix L, Figure 18), the V1 disparity lies below the spurious-torus bulk, whereas mouse EEG overlaps it, reinforcing the distinction between topological recovery and geometric quality.

**Table 3:** Window-level Betti classes, primate V1. % of 200 windows; mean *±* SEM (5,000 block bootstraps).

| Condition | (1, 2, 1) | (1, 1, 1) | Other |
|---|---|---|---|
| Real | 46.5 ± 4.3 | 39.5 ± 5.2 | 14.0 ± 2.0 |
| Fourier shuffle | 33.5 ± 4.7 | 50.0 ± 6.9 | 16.5 ± 4.4 |
| IAAFT | 39.0 ± 4.3 | 29.0 ± 3.3 | 32.0 ± 4.2 |
| Aperiodic 1/*f* | 23.0 ± 3.4 | 47.0 ± 2.7 | 30.0 ± 3.6 |
| Env. phase reset | 32.0 ± 4.4 | 52.0 ± 6.8 | 16.0 ± 3.7 |
| One-mode abl. | 36.5 ± 2.5 | 34.5 ± 4.2 | 29.0 ± 3.9 |

A point cloud can retain the expected loops and enclosed region reflected in Betti numbers even when it is stretched or noisy. Thus, recovering the expected Betti numbers does not guarantee a geometrically clean torus. We therefore further validated the geometric fit of the embedded point cloud to our proposed K-torus with Procrustes disparity, torus score, torus error and *R*^2^. In mouse EEG, the expetec (1, 2, 1) signature appeared in 68.5% of the windows, although its Procrustes disparity was higher than in primate V1 LFP (0.4092 vs. 0.1477), consistent with a less clean embedding. At the population level, torus scores and *R*^2^ values for real recordings are significantly higher than a phase-shuffled null distribution (*p <* 0.001 for all metrics; Figure 5d), while torus error is correspondingly lower. The This separation is stable across a broad range of embedding delays *τ* and window sizes, confirming that the recovered geometry reflects genuine oscillatory structure rather than statistical artifact. Torus-fitting calibration on synthetic AR(2) and AR(4) signals with known ground-truth topology confirms the procedure is not a fitting arteifact (Figure 5c).

## 4 Downstream applications of recovered geometry

Having validated the recovered geometry, we next ask whether it provides useful features for downstream applications. The listed behavior decoding analysis using recovered geometries of neural signals in this section illustrate nominal applications while the method inherently may support other downstream applications. Table 4 summarizes decoding results across three different dataset.

**Table 4:** Decoding performance across recording modalities. Mean SD ±F1 scores. Relevant bands are theta (mouse hippocampal LFP), beta (macaque motor-cortex LFP), and delta (mouse cortical EEG). The final column gives *p* values versus the relevant band and average PSD, respectively.

| Dataset | Torus F1 | Relevant-band F1 | Avg. PSD F1 | *p* (band / PSD) |
|---|---|---|---|---|
| Mouse hippocampal LFP | 0.62 ± 0.10 | 0.59 ± 0.12 | 0.53 ± 0.11 | 0.028 / 3.5 × 10<sup>−7</sup> |
| Macaque motor-cortex LFP | 0.18 ± 0.03 | 0.16 ± 0.03 | 0.16 ± 0.03 | 2.3 × 10<sup>−15</sup> / 3.0 × 10<sup>−13</sup> |
| Mouse cortical EEG | 0.67 ± 0.06 | 0.47 ± 0.05 | 0.49 ± 0.06 | 5.7 × 10<sup>−6</sup> / 5.7 × 10<sup>−6</sup> |

**Table 5:** Positioning relative to delay-embedding and neural-manifold work. Abbreviations — DynSys: dynamical systems; Synth: synthetic; Hipp.: hippocampal; mfld: manifold; pred.: prediction; LDS: linear dynamical system; Betti nos.: Betti numbers from persistent homology. Unlike prior delay-embedding work, which treats lag coordinates as generic reconstruction devices, we prove analytically that oscillatory AR poles produce T^*K*^ topology, validate with formal topological invariants, and demonstrate downstream behavioral decoding. Unlike empirical topology methods Gardner et al. [2022], our *K*-torus is derived from spectral content rather than inferred post-hoc from population activity. Unlike switched and decomposed LDS methods Linderman et al. [2016], Mudrik et al. [2024], Chen et al. [2024], our continuous-drift TVAR formulation preserves a well-defined instantaneous pole structure that maps directly to manifold topology.

| Work | Data | Goal | Metric(s) | Topology claim? |
|---|---|---|---|---|
| Takens (1981) Takens [2006] | DynSys | Embedding theorem | – | – |
| Sauer et al. (1991) Sauer et al. [1991] | DynSys | Embedology | – | – |
| Linderman et al. (2017) Linderman et al. [2016] | Spikes | Switched LDS | Log-lik. | – |
| Ostrow et al. (ICML’24) Ostrow et al. [2024] | Synth | Seq. repr. ≈ delay emb. | Pred. err. | – |
| Raut et al. (Nature’25) Raut et al. [2025] | Pupil→brain | Latent mfld + pred. | *R*<sup>2</sup>, *ρ* | – |
| Gonzalez et al. (JCN’24) Gonzalez et al. [2024] | Hipp. LFP | Geom. coords | Task-dep. | – |
| Gardner et al. (Nature’22) Gardner et al. [2022] | Grid cells | Torus topology (empirical) | Betti nos. | 𝕋<sup>2</sup> (empirical) |
| Mudrik et al. Mudrik et al. [2024], Chen et al. [2024] | Spikes/LFP | Decomposed & probabilistic LDS | *R*<sup>2</sup>, log-lik. | – |
| **This work** | **LFP/EEG** | **Osc. ⇒ 𝕋<sup>*K*</sup>**<br>**(analytic + empirical)** | **Betti nos.,**<br>**torus score, F1** | ***K*-torus**<br>**(theory+empirical)** |

### 4.1 sleep-state decoding

We first examine whether torus parameters distinguish Wake, NREM, and REM in mouse cortical EEG. Across 21 recording hours, LDA trained on 15 geometric features from recovered manifold achieves an F1 score of 0.670 ± 0.060, compared with 0.470± 0.050 for the most relevant individual frequency band and 0.485± 0.062 for average spectral power (Table 4; Figure 6a–d). The distributions for three radii for the 2-torus show separability between sleep stages; while the confusion matrix shows strongest separation for NREM. These results show that the recovered geometry is useful beyond visualization: its compact parameters retain information about sleep state. Mouse hippocampal LFP shows a similar pattern for sleep/awake decoding: torus features achieve 0.624 ± 0.099 F1 versus 0.592 ± 0.124 for theta-band power (Table 4).

**Figure 6:**
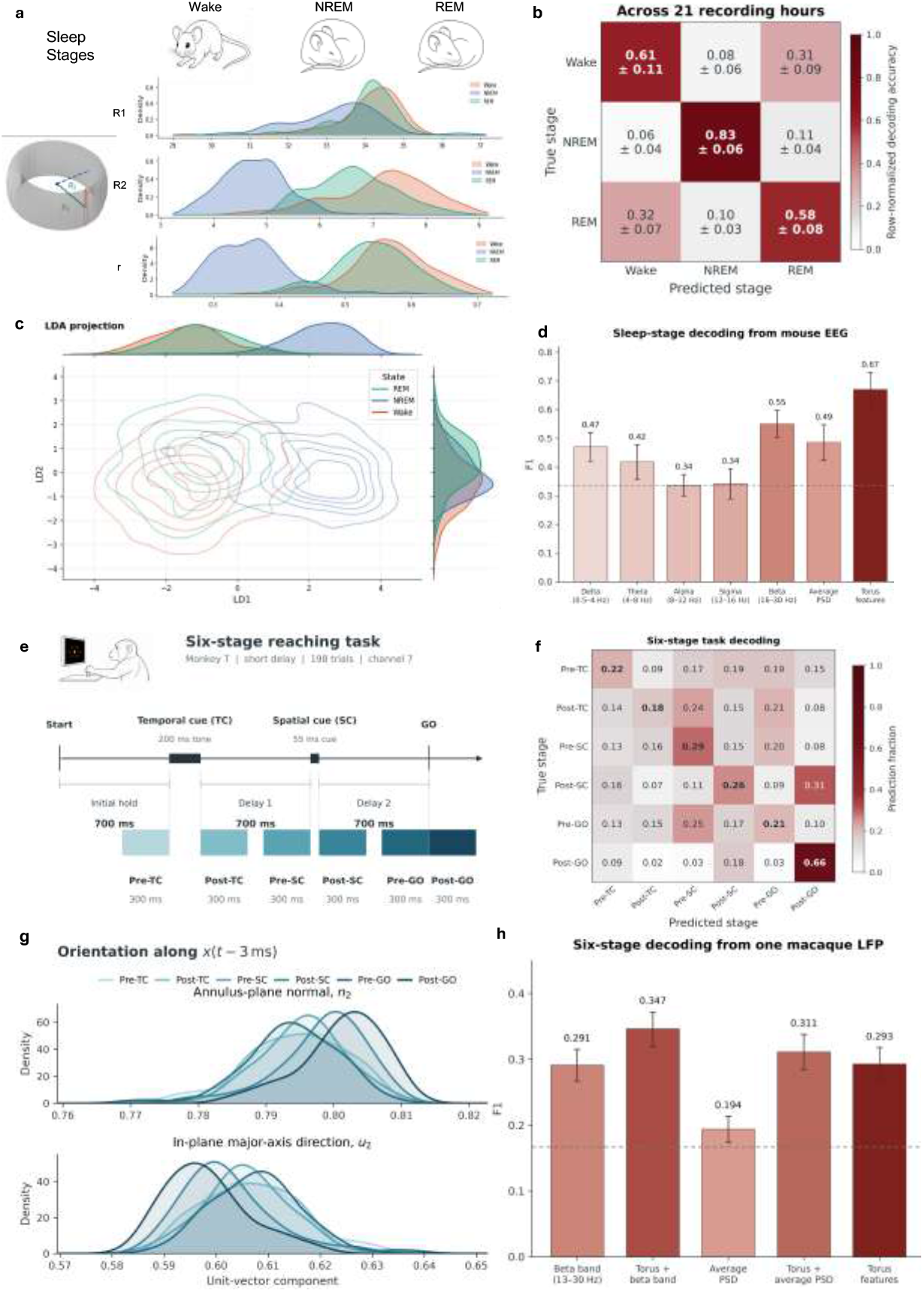
Downstream decoding from lag-embedding geometry. Mouse EEG. **(a–d):** Sleep stages and torus-parameter distributions (a), a confusion matrix across 21 recording hours (b), an LDA projection (c), and F1 scores for torus and spectral features (d). **Macaque LFP (e–h):** The six-stage reaching task and analysis windows (e), stage-decoding confusion matrix (f), stage-dependent orientation features (g), and F1 scores for torus, spectral, and combined features from one LFP (h). Error bars show across-hour standard deviation in (d) and bootstrap confidence intervals in (h).

### 4.2 Macaque behavioral decoding

We next ask whether the same geometric representation carries information about behavior during an active task. We analyzed motor-cortext LFP from two macaques performing the delayed, six-direction reaching task studied by Confais et al. [2020]. The task separates temporal and spatial cues from the eventual movement, allowing us to examine both reach direction and task stage. Across 237 LFP recordings containing all six directions, torus features achieve F1 of 0.181± 0.032 for direction decoding, compared with 0.157 ±0.030 for beta-band power and 0.159± 0.030 for average spectral power (Table 4). We further showed that across animals (macaque n=2), the direction-decoding results are consistent, which are reported in the Appendix N.2.

Because Confais et al. [2020] also found that beta activity varies systematically across task events, we also test whether torus geometry distinguishes task stages. In one Monkey T recording, we decode six 300-ms stages around the temporal cue, spatial cue, and go signal (Figure 6e–h). Torus features achieve 0.293 F1 alone. Adding them to beta-band power raises F1 from 0.291 to 0.347; adding them to average spectral power raises F1 from 0.194 to 0.311 (Appendix N.3). These results provide a single-recording demonstration of task-stage decoding from the recovered geometry.

## 5 Discussion

Our findings establish a theoretical and computational connection between oscillatory neural field dynamics and intrinsic manifold geometry, extending geometric approaches beyond spiking activity. By linking autoregressive dynamics to delay-coordinate topology, we show how independent oscillatory modes can generate predictable toroidal structures. The recovery of these signatures across neural recording modalities, together with their behavioral relevance, suggests that manifold geometry provides not only a descriptive representation of field activity but also a functionally informative characterization of population dynamics.

An important implication concerns the distinction between intrinsic and embedding dimensionality. While conventional embedding representations may improve neural decoding by capturing additional task-related variability, they can also reflect nuisance dimensions or confounding signals. Intrinsic dimensionality, in contrast, characterizes the effective degrees of freedom organizing the dynamics, providing a complementary perspective on their underlying mechanisms Jazayeri and Ostojic [2021], Kong et al. [2026]. Our framework circumvents a longstanding interpretational bias in neural oscillation research, where Fourier-based representations can encourage assumptions about the existence and independence of oscillatory components. By characterizing LFP dynamics without predefined frequency bands or explicit identification of narrowband oscillations, our approach provides a complementary means of investigating independent dynamical degrees of freedom through intrinsic manifold geometry. The toroidal geometry of hippocampal LFPs resembles the continuous-attractor and toroidal structures reported in grid-cell populations Gardner et al. [2022], Chaudhuri et al. [2019]. Although arising from oscillatory dynamics rather than spatial representations, this resemblance motivates direct comparisons between field and spiking manifolds to investigate shared versus distinct principles of neural computation. Such comparisons may further illuminate the relationship between toroidal grid-cell representations Sorscher et al. [2023] and hippocampal dynamics historically associated with theta oscillations Lega et al. [2012], Losonczy et al. [2010], Norman et al. [2025].

Beyond manifold reconstruction, our publicly available TVAR framework opens avenues for investigating slower neural phenomena, including representational drift, cognitive-state transitions, and circuit reorganization, through the evolution of AR coefficients. To facilitate such investigations, we release our implementation on GitHub, providing an open and extensible computational foundation for studying nonstationary neural dynamics across recording modalities and timescales.

## •Technical Appendix •

In the following sections, we provide the theoretical details, implementation choices, synthetic validations, and additional empirical analyses supporting the results in the main paper. The appendix is structured as follows.

## A Related works

### Delay embeddings and state-space reconstruction

Takens’ theorem Takens [2006] and embedology Sauer et al. [1991] establish that delay-coordinate maps embed the underlying state manifold under generic conditions, providing a principled foundation for manifold reconstruction from scalar time series. Recent machine-learning work has revisited this perspective: Ostrow et al. [2024] show that modern sequence models recover hidden state via mechanisms related to delay-coordinate reconstructions; Raut et al. [2025] reconstruct a latent manifold from pupil diameter to predict spatiotemporal brain dynamics; Gonzalez et al. [2024] apply diffusion-mapped delay coordinates to hippocampal LFPs to reveal task-dependent geometric structure. These works treat delay coordinates as generic reconstruction devices and do not provide a principled mapping from oscillatory spectral structure to a specific manifold topology. Our work fills this gap by deriving the topology analytically from the AR pole structure.

### Non-stationary linear dynamical systems for neural data

Switched linear dynamical systems Linderman et al. [2016] and their AR-*p* extensions Lee et al. [2023] model dynamics through discrete latent-state transitions. Decomposed linear systems Mudrik et al. [2024] represent global dynamics as sparse combinations of linear operators, and their probabilistic extensions Chen et al. [2024] improve robustness under dynamical noise. Switched AR-*p* GLMs have been applied to spike trains Escola et al. [2011]. These frameworks model *discrete* regime changes or *globally decomposed* dynamics and therefore do not yield a continuously evolving instantaneous pole structure. Our TVAR formulation with Lipschitz-smooth coefficient drift occupies a complementary niche: it is precisely the regime in which an identifiable T^*K*^ geometry is preserved across time, with a quantified approximation guarantee absent from discrete-switching approaches.

### Topology and geometry as quantitative lenses for neural data

Topology-aware methods have recovered ring structure in head-direction cells via persistent homology Chaudhuri et al. [2019], Giusti et al. [2015], toroidal geometry in grid-cell populations Gardner et al. [2022], and low-dimensional manifold structure in hippocampal dynamics Low et al. [2018], Nieh et al. [2021]. Our work differs fundamentally: the toroidal geometry is not inferred post-hoc from population activity but is *derived analytically* from the LFP’s AR pole structure and then independently confirmed using persistent homology on the recovered embedding — providing both a predictive theory and its topological validation.

### LFP Spectral Structure and Neural Manifold Geometry

Parameterizing LFP/EEG spectra into periodic and aperiodic components Donoghue et al. [2020], Choubdar et al. [2023] has enabled principled cross-condition comparisons beyond fixed frequency bands, and oscillatory LFP features have been linked to attention, memory, and cognitive control Miller et al. [2018], Gedankien et al. [2025], Fazlali et al. [2020], Zabeh et al. [2023]. Our framework provides a geometric reinterpretation of these oscillatory signatures: sustained rhythms are coordinate directions of the LFP manifold, and their geometric parameters carry behaviorally relevant information. This geometric perspective further enables, for the first time, a principled comparison of the topology of spiking and LFP manifolds, providing a unified framework for investigating spike–field interactions and bridging a longstanding gap between these complementary representations of neural population dynamics Kolibius et al. [2026], Guth et al. [2025], Qasim and Kunz [2023].

## B Technical companion for “Analytical Characterization of LFP Manifold Geometry”

This appendix is a self-contained technical companion to Section 2 in the main paper. The main text states the core narrative (oscillations → phase variables → torus under delay embedding), while this appendix provides (i) precise model definitions, (ii) algebraic equivalences (AR↔ companion form), (iii) the minimal-lag embedding result (two coordinates per oscillatory mode), (iv) a PSD-level correspondence between peaks and poles, and (v) the conditioning interpretation used in the equalization discussion.

### Roadmap

Appendix A.1 defines the (multi-channel) AR/VAR model and clarifies the stability/boundedness assumptions. Appendix A.2 gives the companion/state-space form used to interpret delay coordinates as a linear dynamical state. Appendix A.3 proves the geometric core: sustained oscillatory modes imply a *K*-torus under a minimal 2*K*-lag embedding. Appendix A.4 states the AR PSD formula, connects empirical PSD decompositions (peaks + 1*/f*^*α*^ background) to poles, and explains why delay embeddings carry strictly more information than the power spectrum alone. Appendix A.5 formalizes the conditioning discussion: oscillatory modes determine topology, while background/noise affect thickness and anisotropy.

### B.1 Autoregressive model and multichannel reduction

The main text presents a single-channel AR model for clarity. Here we write the multichannel version and show that it reduces to a block-diagonal VAR(*p*).

#### Multichannel process

Let *X*(*t*) = [*x*_1_(*t*), …, *x*_*N*_ (*t*)]^*⊤*^ ∈ ℝ^*N*^ denote an *N* -channel discrete-time process. Assume each channel follows an AR model

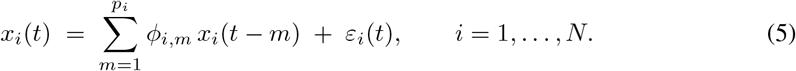

Stacking innovations gives *ε*(*t*) = [*ε*_1_(*t*), …, *ε*_*N*_ (*t*)]^*⊤*^. We assume E[*ε*(*t*)] = 0, temporal whiteness, and allow cross-channel correlation: E[*ε*(*t*)*ε*(*t*)^*⊤*^] = *R*_*ε*_ (constant for simplicity).

#### Stability/boundedness assumption (used throughout)

We assume the AR dynamics are stable (all poles strictly inside the unit disk) except for a subset of *oscillatory factors* that may lie (or be arbitrarily close) to the unit circle and are treated as the sustained modes of interest. This assumption is consistent with analyzing finite windows of locally stationary data and prevents pathological growth. (When oscillatory poles are exactly on the unit circle, statements below should be read in the noiseless/asymptotic idealization.)

##### Lemma B.1

(Block–diagonal AR(*p*) reduction). *Let p* := max_*i*_ *p*_*i*_ *and set ϕ*_*i,m*_ := 0 *for all m > p*_*i*_ *(zero padding). Define diagonal lag blocks*

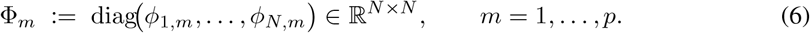

*Then the stacked process obeys the block–diagonal VAR(p)*

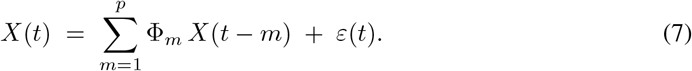

*Proof*. The *i*th entry of 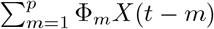 equals 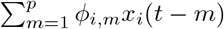, which by zero padding reduces to 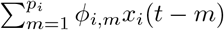. Stacking over *i* yields (7).

#### Subsequent main text rational

The main text analyzes a single channel for readability; Lemma B.1 shows that multichannel extensions follow by stacking, and do not change the oscillation geometry mechanism.

### B.2 Companion form: AR dynamics as a linear state recursion

Delay embeddings are naturally expressed by stacking lagged samples. The companion form makes this stacking explicit as a linear dynamical state.

#### Lemma B.2

(Companion form and equivalence to VAR(*p*)). *Let X*(*t*) *satisfy* (7). *Define the stacked lag state*

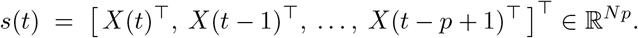

*Define the companion matrices*

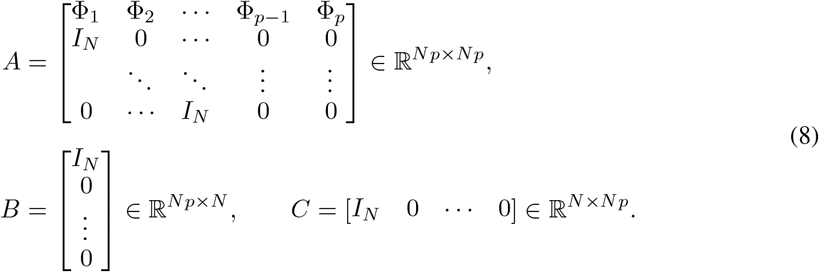

*Then*

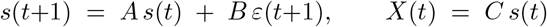

*is algebraically equivalent to* (7). *Moreover*,

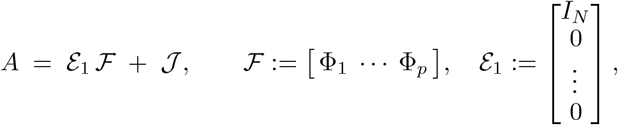

*where J is the fixed lag-shift matrix, so all learnable dynamics reside in* ℱ.

*Proof*. The first block row yields 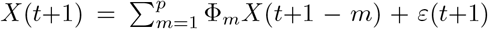, i.e. (7). The remaining block rows implement the lag shift. The decomposition follows by inspection.

#### Connection to delay embeddings

Any delay embedding using *p* lags is a linear projection of the companion state *s*(*t*). Thus the geometry of delay embeddings is governed by the oscillatory content of the AR state.

### B.3 Oscillatory poles → phases → a *K*-torus

This section formalizes the geometric claim in the main text: *each sustained oscillatory mode contributes one phase variable on S*^1^; *K modes yield* T^*K*^.

#### B.3.1 Factorization into oscillatory and stable components

For channel *i* define the characteristic polynomial

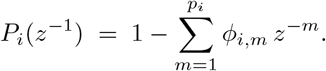

##### Lemma B.3

(Pole factorization). *For each channel i there exist an integer K*_*i*_ ≥ 0 *and frequencies* 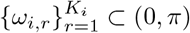 *such that*

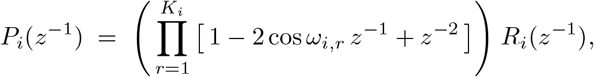

*where all roots of the residual factor R*_*i*_(*z*^*−*1^) *lie strictly inside the open unit disk*.

*Proof*. Factor *P*_*i*_ over ℂ as (1 *ζz*^*−*1^). Real coefficients imply conjugate pairing. Each unit-modulus conjugate pair *ζ* = *e*^*iω*^, 0 *< ω < π*, gives

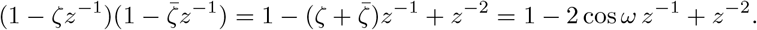

All remaining roots are strictly stable by assumption, forming *R*_*i*_.

##### Meaning

Each quadratic factor corresponds to one sustained oscillatory mode (one independent phase). The residual stable factor corresponds to non-oscillatory components that affect *conditioning/thickness* but not topology.

#### B.3.2 Minimal lags: two coordinates per oscillation

##### Lemma B.4

(Minimal-lag linearization and invertibility). *Suppose channel i has K*_*i*_ *sustained modes with distinct frequencies modulo π. Consider the noiseless oscillatory component*

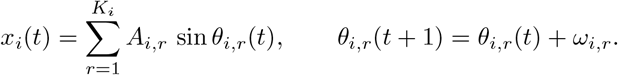

*Stack* 2*K*_*i*_ *consecutive lag* 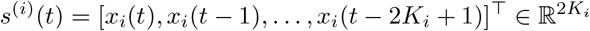. *Then there exist an invertible* 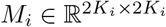, *a positive diagonal D*_*i*_, *and*

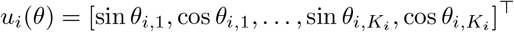

*such that*

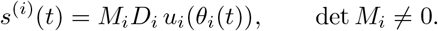

*Proof*. Using sin(*α* − *β*) = sin *α* cos *β* − cos *α* sin *β*, each lag *x*_*i*_(*t* − *k*) is linear in (sin *θ*_*i,r*_(*t*), cos *θ*_*i,r*_(*t*)) with coefficients cos(*kω*_*i,r*_) and − sin(*kω*_*i,r*_). Stacking *k* = 0, …, 2*K*_*i*_ − 1 yields *s*^(*i*)^(*t*) = *M*_*i*_*D*_*i*_*u*_*i*_(*θ*_*i*_(*t*)). Distinctness modulo *π* implies full rank of the resulting realified Vandermonde-type matrix *M*_*i*_.

##### This is the formal version of “two lags per oscillation.”

Each oscillatory mode carries one phase *θ* but requires two real coordinates (sin *θ*, cos *θ*). Lemma B.4 shows that 2*K* delay coordinates are sufficient (and generically necessary) to obtain an embedding.

#### B.3.3 Global *K*-torus embedding

Stacking across channels yields a 2*K*-dimensional minimal lag vector *s*_min_(*t*). Define *u*(*θ*) = [sin *θ*_1_, cos *θ*_1_, …, sin *θ*_*K*_, cos *θ*_*K*_]^*⊤*^ and let *M, D* be the block-diagonal stacks of *M*_*i*_, *D*_*i*_.

##### Proposition B.5

(*K*-torus embedding). *Assume the hypotheses of Lemma B*.*4 hold so that M is invertible. Define F* : T^*K*^ → ℝ^2*K*^ *by*

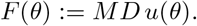

*Then F is a smooth embedding and ℳ*:= *F* (T^*K*^) *is a compact manifold diffeomorphic to* (*S*^1^)^*K*^. *Equivalently, in whitened coordinates y* = (*MD*)^*−*1^*s, the manifold satisfies*

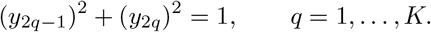

*Proof. u* is smooth and *MD* is invertible, so *F* is smooth. D*u* has rank *K* everywhere and thus D*F* = *MD* D*u* has rank *K* everywhere (immersion). Injectivity follows from *F* (*θ*) = *F* (*θ*^*′*^)⇒ *u*(*θ*) = *u*(*θ*^*′*^) after applying (*MD*)^*−*1^, which implies equality of each (sin *θ*_*q*_, cos *θ*_*q*_) pair and hence 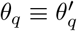 (mod 2*π*).

### B.4 PSD-level correspondence: peaks ↔ poles

This section supports the main-text claim that spectral peaks identify sustained oscillatory modes, and explains why delay embeddings carry strictly more geometric information than the power spectrum alone.

#### AR PSD

For a (single-channel) AR(*p*) process

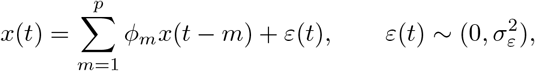

the PSD is

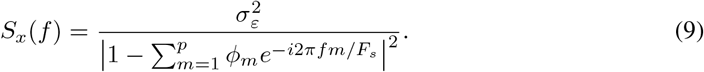

If the characteristic polynomial has a conjugate pole pair *z* = *ρe*^*±iω*^ with *ρ* ≈1, then (9) exhibits a narrowband peak near frequency *ω*. Stable poles with *ρ <* 1 broaden the spectrum and contribute to the aperiodic floor.

#### Composite PSDs used in neuroscience

A common empirical decomposition is

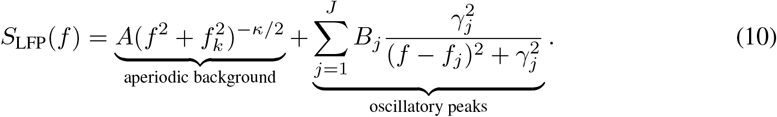

The AR viewpoint corresponds to realizing each oscillatory peak with an AR(2) factor having conjugate poles near the unit circle, and realizing the aperiodic term with strictly stable (typically real) poles that shape slope/offset over the band of interest. This gives the correspondence used in the main text:

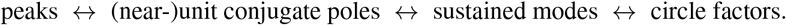

#### Why delay embeddings carry more information than the PSD

The correspondence above is *one-way lossy* for the power spectrum: because 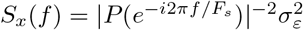 depends only on the moduli of the pole transfer function evaluated on the unit circle, it discards all phase information. Concretely, two AR processes with the same pole moduli {*ρ*_*r*_} and frequencies {*ω*_*r*_} but different initial phase offsets {*φ*_*r*_} produce identical PSDs yet trace out different trajectories in delay-coordinate space. When two oscillatory modes are nearby in frequency 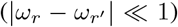, their spectral peaks may overlap and become indistinguishable in the PSD, while the delay embedding continues to resolve them as two independent circular factors provided their phases are not locked. More formally, Proposition B.5 establishes that the topology of the delay embedding is determined by the number of *distinct* sustained pole pairs modulo *π*, a condition that is invisible to the PSD whenever modes are spectrally proximate. Delay embeddings therefore resolve phase-structure ambiguities that the power spectrum cannot, recovering the underlying T^*K*^ geometry even when the PSD is uninformative about the true value of *K*.

### B.5 Conditioning: topology vs. thickness (equalization)

The main text notes that linear spectral equalization improves the empirical recoverability of toroidal structure without altering the underlying topology. This section provides the algebraic basis for that claim by formalizing how oscillatory amplitudes, aperiodic background, and noise affect the geometric conditioning of the delay embedding while leaving *K* — and hence the Betti numbers — unchanged.

In minimal-lag form, the noiseless embedding satisfies

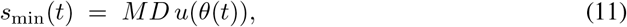

where *D* = diag(*A*_1_, *A*_1_, …, *A*_*K*_, *A*_*K*_) encodes oscillation amplitudes and *M* is the invertible realified Vandermonde map from Lemma B.4. The image of T^*K*^ under *F* (*θ*) = *MD u*(*θ*) is thus an *anisotropically scaled* torus: circle factor *q* has effective radius *A*_*q*_ ·*σ*_*q*_, where *σ*_*q*_ denotes the corresponding singular value of *M*. When amplitudes {*A*_*k*_} are strongly imbalanced, weaker oscillatory modes are embedded at much smaller geometric scale and become difficult to resolve against measurement noise or in low-dimensional projections.

#### Equalization as conditioning improvement

Spectral equalization applies an invertible diagonal rescaling Λ = diag(*λ*_1_, *λ*_1_, …, *λ*_*K*_, *λ*_*K*_) to the embedding coordinates, replacing *MD* by *MD*Λ^*−*1^. Choosing *λ*_*k*_ = *A*_*k*_ (or any estimate thereof) normalizes the amplitude factors in *D*, improving the condition number of *MD* without altering the rank or the topology of ℳ: the manifold remains diffeomorphic to (*S*^1^)^*K*^ by Proposition B.5, since Λ^*−*1^ is invertible. Thus equalization *reveals* existing toroidal structure rather than creating new geometry, consistent with the interpretation given in the main text.

#### Role of aperiodic background and noise

Additive aperiodic components and measurement noise enter the embedding as

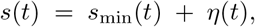

where *η*(*t*) collects contributions from stable AR poles and innovation noise. Because stable poles produce no unit-circle factors (Lemma B.3), *η*(*t*) introduces no additional circular degrees of freedom. Geometrically, *η*(*t*) inflates the point cloud around the torus skeleton ℳ, increasing its effective thickness without changing *K*. The signal-to-noise ratio for detecting the *k*-th circular factor is approximately *A*_*k*_*/* ∥*η* ∥; equalization maximizes the minimum SNR across factors, making the full T^*K*^ structure uniformly detectable. A detailed simulation study of how the spectral exponent *α* of the aperiodic background affects embedding thickness and conditioning is provided in Appendix J.

#### Takeaway

Oscillatory modes determine *K* (topology); aperiodic structure and noise determine the thickness and anisotropy (geometric conditioning) of the observed point cloud around the underlying torus. Equalization addresses conditioning without touching topology, and its effect is largest when oscillatory amplitudes are imbalanced across modes.

## C Local Stationarity Bridge — Full Derivation and Empirical Validation

This appendix provides the full technical details supporting the local stationarity bridge in Section 3. We first derive the bound on the discrepancy between the TVAR dynamics and the frozen AR surrogate (Appendix C.1), then state the formal result on polynomial root continuity (Appendix C.2), and finally validate Condition (4) empirically across all biological datasets used in this work (Appendix C.3).

### C.1 Full Derivation of the Local Approximation Bound

We model the LFP signal as a time-varying autoregressive (TVAR) process,

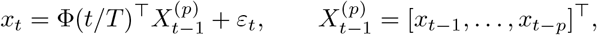

where the coefficient path Φ : [0, 1] → ℝ^*p*^ is Lipschitz with constant *L*_Φ_, i.e.,

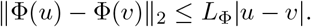

Fix a reference time *t*_0_, and define the frozen-coefficient approximation

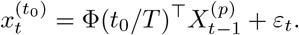

The pointwise discrepancy between the TVAR process and this surrogate is

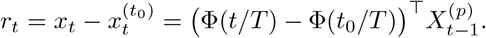

We now bound |*r*_*t*_| over a local window of half-width *w* centered at *t*_0_. Since the LFP signal is bounded in amplitude as |*x*_*t*_| ≤ *B* over the window, the lag vector satisfies

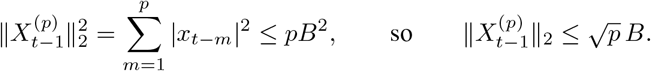

Applying the Cauchy–Schwarz inequality and the Lipschitz condition, for any |*t* − *t*_0_| ≤ *w*,

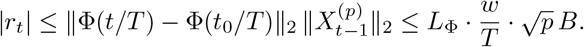

Therefore, whenever

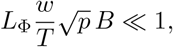

the TVAR process is well approximated on the local window by its frozen AR surrogate, and the approximation error vanishes as *w/T*→ 0 for fixed *L*_Φ_, *p*, and *B*. This confirms that working in short locally stationary windows is sufficient to reduce the TVAR dynamics to a fixed-coefficient AR model up to a controllable error.

#### Remark C.1.

The bound scales as *O*(*w/T*), confirming that shorter windows yield better local approximations. In practice, *w* must be chosen to balance this approximation error against estimation variance: windows that are too short yield noisy AR coefficient estimates, while windows that are too long violate local stationarity. We discuss this tradeoff and our window length selection procedure in Appendix F.

### C.2 Polynomial Root Continuity

The bound in Appendix C.1 controls the discrepancy in AR coefficients between the TVAR process and its frozen surrogate. To conclude that the lag-embedded geometry is also approximately preserved, we need that small coefficient perturbations imply small perturbations to the pole configuration. This follows from the classical fact that polynomial roots are continuous functions of their coefficients.

#### Proposition C.2

(Continuity of polynomial roots). *Let* 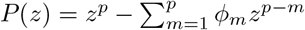 *be the characteristic polynomial of an AR*(*p*) *process with coefficients* ***ϕ*** = [*ϕ*_1_, …, *ϕ*_*p*_]^*⊤*^. *For any ϵ >* 0, *there exists δ >* 0 *such that if* ∥***ϕ***− ***ϕ***^*′*^ ∥_2_ *< δ, then the roots of P and P*^*′*^ *can be matched so that each pair differs by at most ϵ in modulus and argument*.

This result is classical and follows from Rouché’s theorem; see Marden [1966] for a detailed treatment. A direct consequence is that whenever 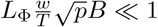, the poles of the frozen surrogate lie within an *O* (*L*_Φ_*w/T*) neighborhood of the instantaneous TVAR poles. In particular, oscillatory poles that are sustained (near-unit-modulus) in the TVAR process remain near-unit-modulus in the surrogate, and the number of sustained oscillatory modes *K* is preserved. By Proposition 2.1, the lag-embedded geometry therefore tracks a *K*-torus determined by the instantaneous oscillatory modes up to a small geometric perturbation.

### C.3 Empirical Validation of the Local Stationarity Condition

Condition (4) makes a concrete, testable prediction: the product 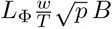 should remain well below 1 across the analysis windows used in our experiments. We verify this directly on all three biological datasets.

#### Estimation procedure

For each recording session and channel, we estimate *L*_Φ_ from finite differences of OLS-fitted AR coefficients across adjacent non-overlapping windows of length *w*:

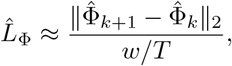

where 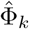 denotes the OLS coefficient estimate in window *k*. The amplitude bound *B* is estimated as the 99th percentile of |*x*_*t*_| within each window. The embedding order *p* is set by the elbow criterion on held-out prediction error, as described in the main text.

#### Discussion

Across all recording sessions, channels, and window lengths considered, the condition 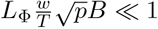 holds comfortably (Table 6, Figure 7), confirming that the locally stationary regime is a valid operating assumption for the LFP signals analyzed in this work rather than a purely theoretical idealization. These results justify applying Proposition 2.1 within each analysis window and ground the time-varying reconstruction framework of Section 3 in empirically verified assumptions.

**Table 6:**
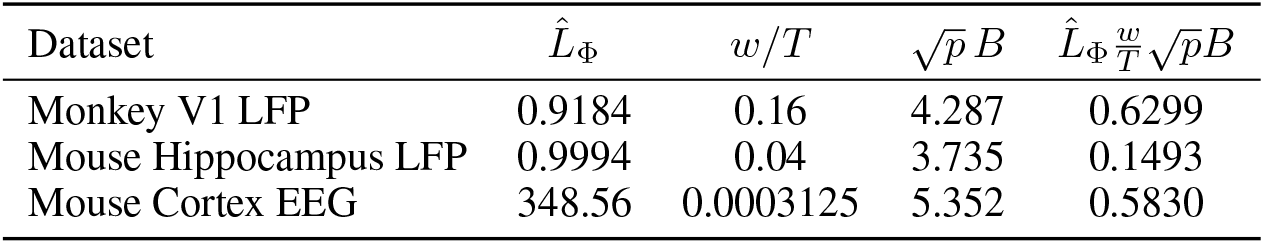
Empirical evaluation of the local stationarity condition across datasets. Values report mean ±standard deviation across recording sessions and channels. The condition 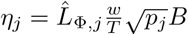 is satisfied comfortably in all cases.

| Dataset | $\hat{L}_{\Phi}$ | $w/T$ | $\sqrt{p}\,B$ | $\hat{L}_{\Phi}\frac{w}{T}\sqrt{p}B$ |
|---|---|---|---|---|
| Monkey V1 LFP | 0.9184 | 0.16 | 4.287 | 0.6299 |
| Mouse Hippocampus LFP | 0.9994 | 0.04 | 3.735 | 0.1493 |
| Mouse Cortex EEG | 348.56 | 0.0003125 | 5.352 | 0.5830 |

**Figure 7:**
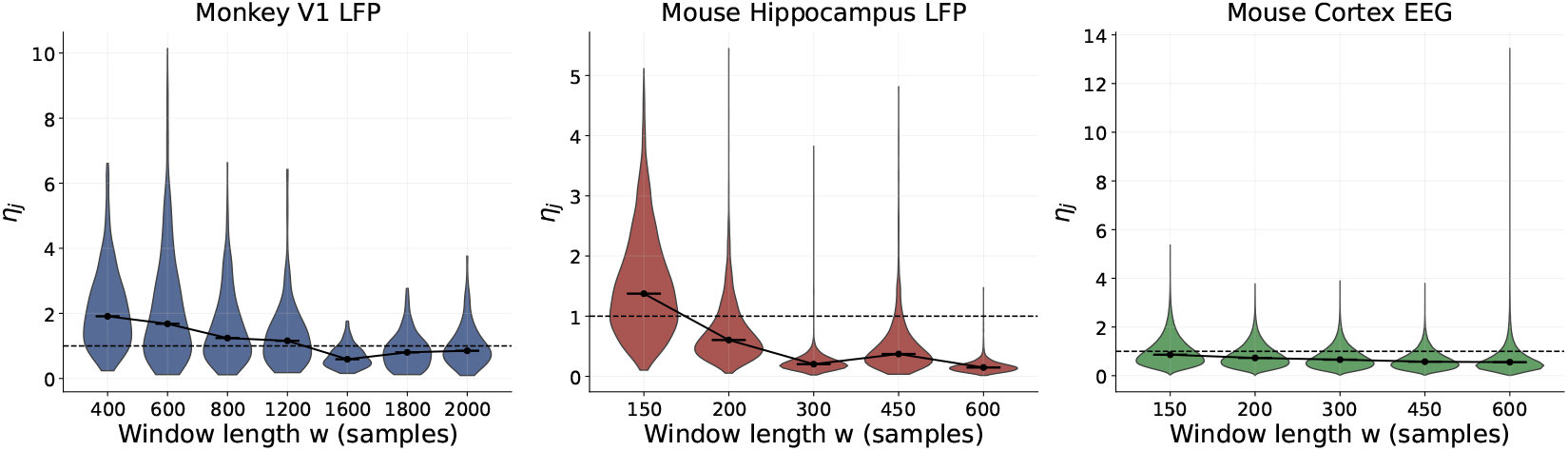
Distribution of the local-stationarity score across window lengths for each dataset. Each violin shows the window-level distribution of 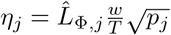 at a fixed window length *w*, with the median marked in black. The dashed horizontal line indicates the reference threshold *η* = 1.

Furthermore, the estimated 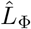 values are consistent with the smoothness regularization imposed by ℒ_smooth_ in DeepLagField: by penalizing rapid temporal variation in the learned coefficients, the architecture directly enforces the Lipschitz condition that underlies this analysis, providing a principled connection between the theoretical assumption and the model design.

## D Additional Analysis Results on TVAR Function Classes

This appendix provides a more granular evaluation of **DeepLagField** across diverse classes of time-varying autoregressive (TVAR) coefficient functions. The goal is to assess robustness beyond aggregate benchmarks by probing how coefficient dynamics with different smoothness, periodicity, and nonlinearity affect reconstruction accuracy and order recovery.

### D.1 Benchmark Setup

We evaluate TVAR coefficient estimation on eight representative function classes: *sinusoidal, Fourier series, quasi-periodic, polynomial drift, logistic transition, multi-sigmoid, Gaussian bumps*, and *smooth random*. Each class induces distinct temporal structure ranging from smooth periodic variation to abrupt but continuous transitions.

For each class, we generate multiple realizations with ground-truth order *p* ∈ {2, 3, 4, 5, 6} and report results averaged across orders and seeds. Performance metrics include: (i) coefficient mean-squared error, (ii) signal reconstruction error, and (iii) order deviation statistics.

### D.2 Quantitative Results Across Function Classes

Table 7 summarizes performance of **DeepLagField** across all coefficient families. Results indicate stable reconstruction accuracy and consistent order estimation across a wide range of temporal behaviors.

**Table 7:** Performance of DeepLagField across TVAR coefficient function classes. Values report mean performance averaged over orders *p* ∈ {2, …, 6} and seeds. Lower is better (↓), higher is better (↑).

| Function class | Coeff. MSE ↓ | Signal MSE ↓ | Δ*p* ↓ | Δ*p*/*p* ↓ |
|---|---|---|---|---|
| Sinusoidal | 0.003924 | 0.183046 | 1.085483 | 0.290002 |
| Fourier series | 0.004272 | 0.184083 | 1.069846 | 0.284058 |
| Quasi-periodic | 0.004035 | 0.182153 | 1.063116 | 0.281938 |
| Polynomial drift | 0.004315 | 0.183816 | 1.109580 | 0.302322 |
| Logistic transition | 0.003673 | 0.183003 | 1.106807 | 0.300061 |
| Multi-sigmoid | 0.003155 | 0.182505 | 1.132916 | 0.309419 |
| Gaussian bumps | 0.003542 | 0.183888 | 1.123097 | 0.302140 |
| Smooth random | 0.007759 | 0.189260 | 1.221906 | 0.343524 |

**Table 8:**
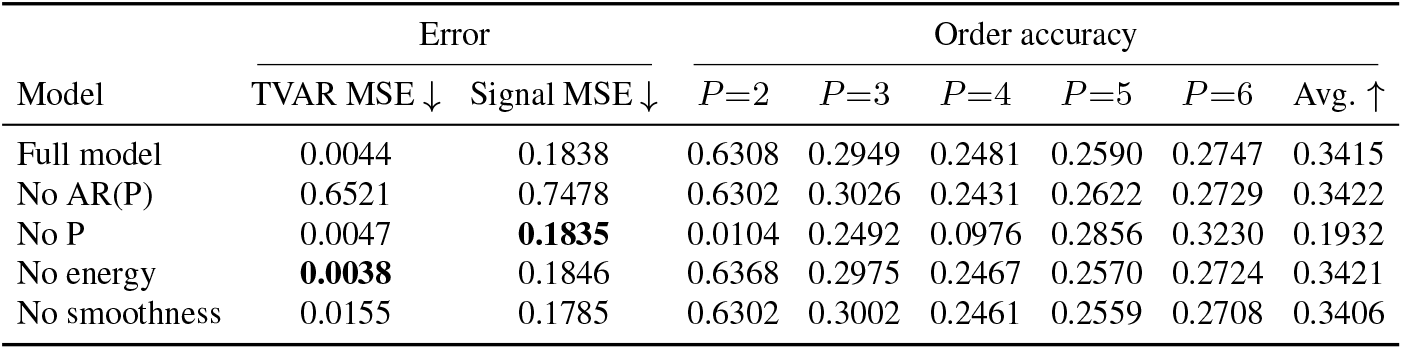
Full ablation results on time-varying AR benchmarks. While several ablations preserve low prediction error, stable recovery of autoregressive order degrades substantially when key structural constraints are removed.

|  | Error |  | Order accuracy |  |  |  |  |  |
|---|---|---|---|---|---|---|---|---|
| Model | TVAR MSE ↓ | Signal MSE ↓ | *P*=2 | *P*=3 | *P*=4 | *P*=5 | *P*=6 | Avg. ↑ |
| Full model | 0.0044 | 0.1838 | 0.6308 | 0.2949 | 0.2481 | 0.2590 | 0.2747 | 0.3415 |
| No AR(P) | 0.6521 | 0.7478 | 0.6302 | 0.3026 | 0.2431 | 0.2622 | 0.2729 | 0.3422 |
| No P | 0.0047 | **0.1835** | 0.0104 | 0.2492 | 0.0976 | 0.2856 | 0.3230 | 0.1932 |
| No energy | **0.0038** | 0.1846 | 0.6368 | 0.2975 | 0.2467 | 0.2570 | 0.2724 | 0.3421 |
| No smoothness | 0.0155 | 0.1785 | 0.6302 | 0.3002 | 0.2461 | 0.2559 | 0.2708 | 0.3406 |

**Table 9:** Scalability analysis of DeepLagField across hidden dimensions. Experiments were conducted on an NVIDIA RTX 5090 GPU with 32GB memory and an AMD Threadripper PRO 9975WX CPU. Timing was averaged over 200 steps with batch size 32 and sequence length *T* = 600. Over the evaluated range, approximately 160K–3M parameters, training and inference times remain nearly constant, although computational cost is expected to increase for substantially larger configurations.

| *d* | Params | Training<br>Compute (ms) | Inference<br>Compute (ms) | Train<br>(samples/s) | Infer<br>(samples/s) |
|---|---|---|---|---|---|
| 32 | 159.7K | 0.80 ± 0.01 | 0.30 ± 0.00 | 39,885 | 105,762 |
| 64 | 319.9K | 0.81 ± 0.00 | 0.31 ± 0.00 | 39,268 | 102,175 |
| 128 | 652.6K | 0.83 ± 0.01 | 0.30 ± 0.00 | 38,690 | 104,967 |
| 256 | 1.37M | 0.85 ± 0.00 | 0.31 ± 0.00 | 37,467 | 104,485 |
| 512 | 2.99M | 0.85 ± 0.00 | 0.32 ± 0.00 | 37,863 | 101,538 |

#### Discussion

Performance is strongest for smooth oscillatory and quasi-periodic coefficients, and degrades gracefully for more irregular but continuous dynamics. Notably, no function class leads to catastrophic failure, indicating that the physics-informed regularization stabilizes learning across regimes.

### D.3 Order Recovery Confusion Analysis

To further understand order estimation behavior, we report confusion matrices for predicted versus true AR order across function classes (Figure 8). Each matrix aggregates predictions across all seeds and realizations within a class.

**Figure 8:**
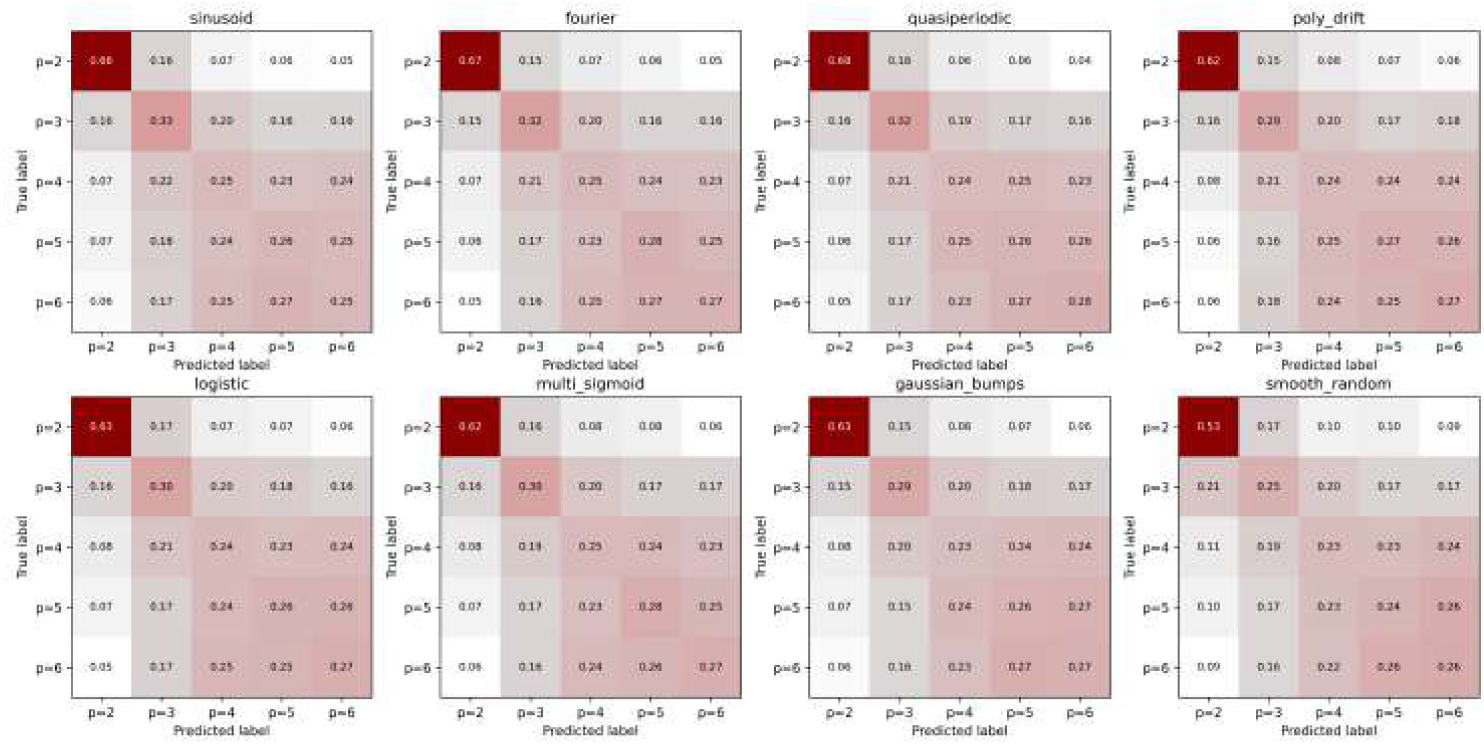
Order recovery confusion matrices across TVAR coefficient classes. Rows correspond to true AR order (*p* = 2–6) and columns to predicted order. While exact order recovery is challenging in highly nonstationary settings, predictions remain concentrated near the diagonal, indicating bounded error and absence of systematic bias. Errors are typically ± 1 and increase mildly for smoother, slowly varying coefficient functions where multiple orders provide a similar predictive fit.

#### Interpretation

Order ambiguity arises primarily from intrinsic identifiability limits: different AR orders can explain the same local dynamics equally well under noise. Importantly, the confusion matrices show no collapse to trivial solutions and no tendency to overestimate order, supporting the stability of the learned lag structure.

#### Takeaway

These additional results demonstrate that **DeepLagField** maintains consistent reconstruction accuracy and interpretable order estimates across a broad family of TVAR dynamics. This robustness supports its use as a general-purpose tool for analyzing locally stationary oscillatory signals beyond synthetic benchmarks.

### D.4 Model Coefficient Recovery Analysis

To understand the ability of *DeepLagField* on coefficient recovery, we report results of *DeepLagField*’s predictions on coefficients from sample families of the benchmark dataset (Figure 9). We further investigate the residual errors of coefficient predictions of trained and untrained DeepLagField (Figure 10).

**Figure 9:**
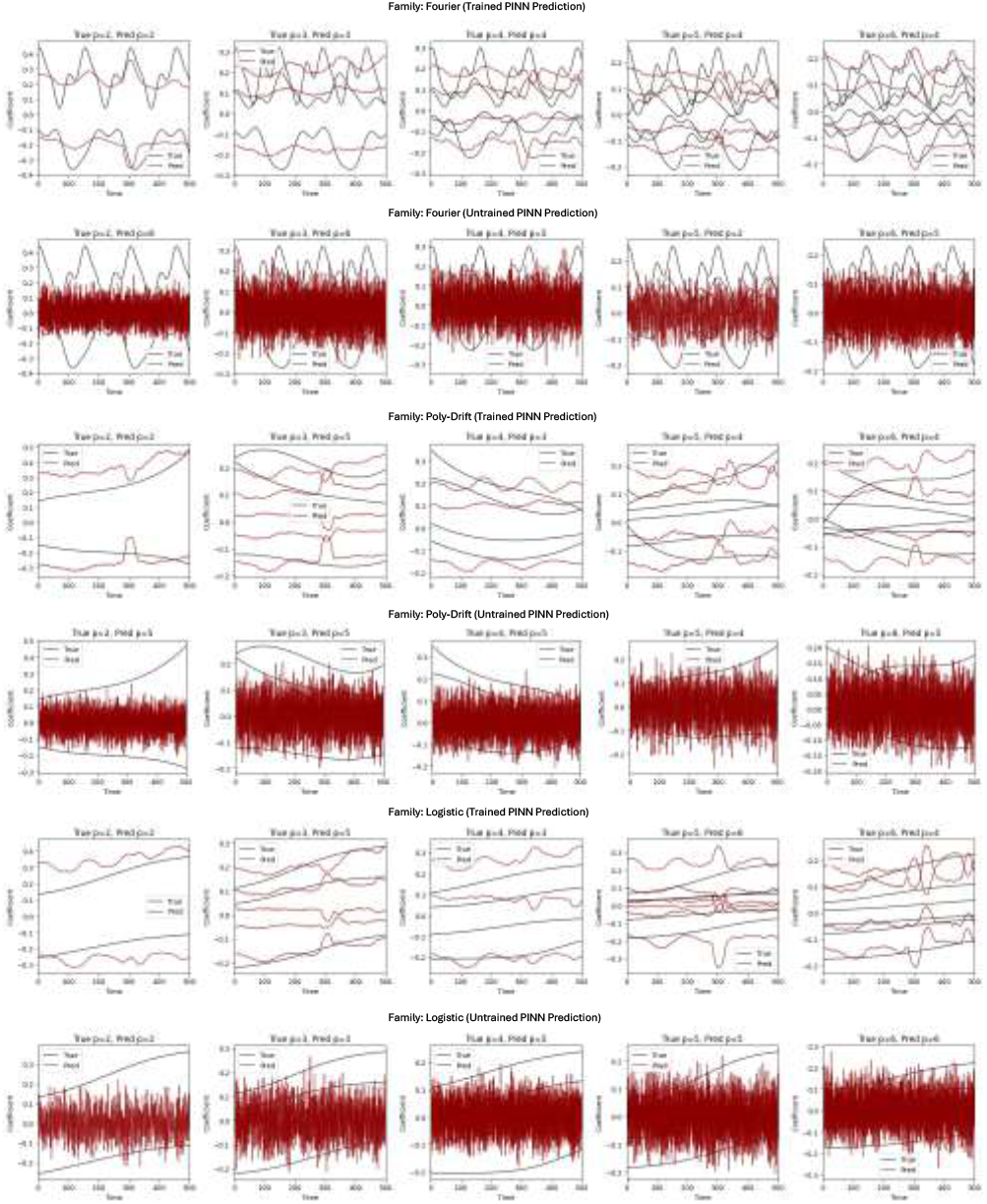
Examples of coefficient prediction by trained vs. untrained PINN on time-varying AR benchmark. Each pair of rows shows predictions from the trained model (rows 1, 3, 5) and an untrained model with random weights (rows 2, 4, 6) for the Fourier, Polynomial-Drift, and Logistic signal families respectively. The trained model accurately captures both the dynamics and smoothness of the time-varying coefficients, whereas the untrained model produces random, jagged predictions — indicating that prediction accuracy stems from learning rather than the PINN architecture alone.

**Figure 10:**
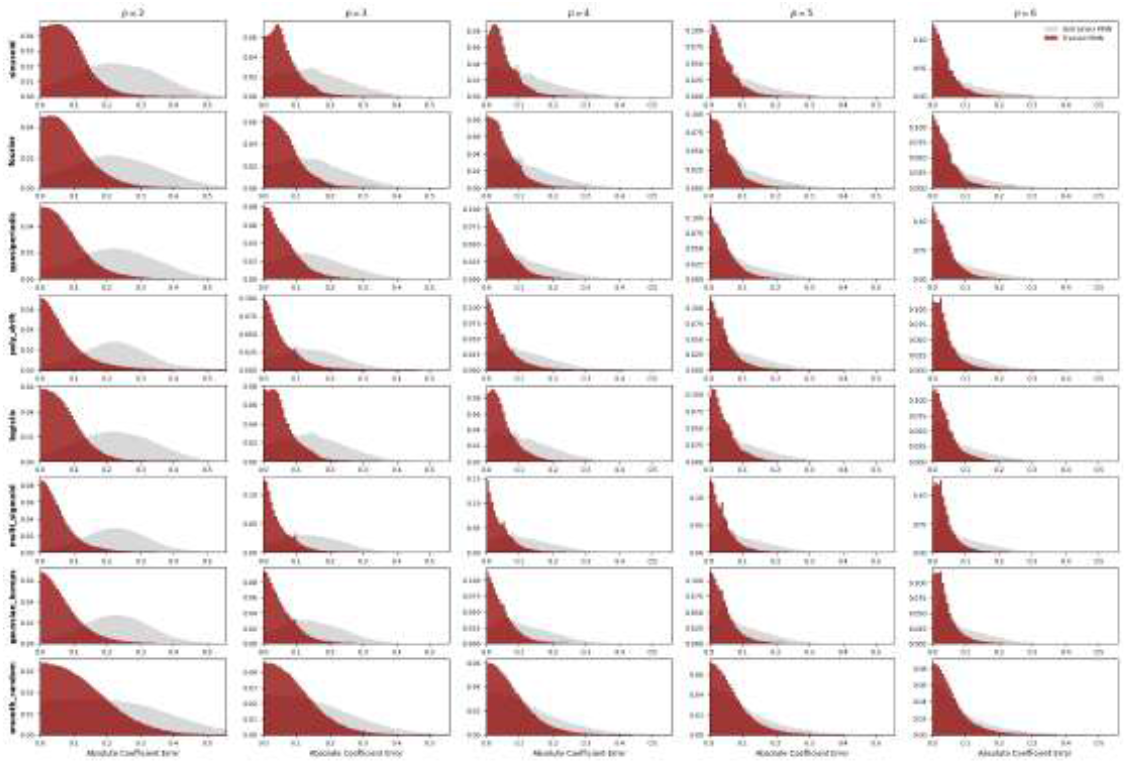
Examples of coefficient prediction by trained vs. untrained PINN on time-varying AR benchmark. Distribution of coefficient prediction error across benchmark dataset consisting of 8 signal families (rows) and AR orders spanning 2 – 6 (columns). Each panel shows overlapping histograms for an untrained PINN model with random weights (gray) and the trained PINN (dark red). The distributions of trained PINN error hug around 0 with a decreasing thin tail from order 2 to 6 in all families, compared with the untrained PINN error baselines showing a random more spread-out errors. The overall trained PINN prediction errors consistently near 0 shows that PINN is robust in predicting both different families of signal and different order of AR signal, which shows a consistent trend with the examples in figure 9.

### D.5 Training Stability of *DeepLagField*

To assess the training stability of *DeepLagField* on benchmark dataset, figure 11 shows the total training and validation losses and oder prediction accuracy over 100 epochs. Both losses decrease monotonically and stabilize after 40 epochs, while both training and validation order accuracy stabilize after 20 epochs. We note that the validation loss and order accuracy suggest that more extensive training would likely show an improvement in model performance.

**Figure 11:**
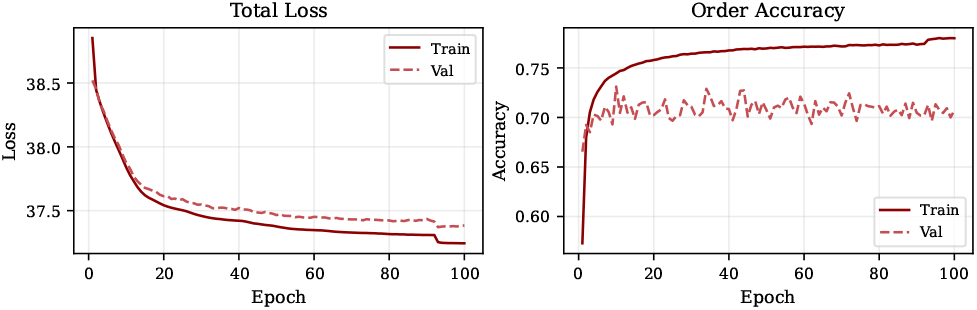
Training and validation curves for total loss and order accuracy DeepLagField on benchmarks. Training (solid line) and validation (dashed line) total loss and order accuracy (defined as the fraction of samples for which 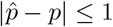, where *p* is the ground-truth AR order and 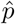 is the predicted order) show convergence, showing training stability.

## E Additional Empirical Results on Real Neural Recordings

This appendix presents additional empirical results on real neural recordings that support the main claims of the paper. The objectives are threefold: (i) to demonstrate robustness of the recovered geometry across recording channels, time windows, and species; (ii) to illustrate how the inferred geometry is expressed locally in time in nonstationary signals; and (iii) to document representative success and limitation cases, clarifying the scope of the proposed framework. All analyses follow exactly the same pipeline as in the main text, with no dataset-specific tuning.

### E.1 Manifold Visualization for Mouse Recordings

Figures 12 and 13 extend the manifold reconstruction pipeline of Figure 3 in the main text to two independent mouse recording modalities: hippocampal local field potential (LFP) and cortical electroencephalography (EEG). These results were omitted from the main manuscript due to space constraints; they are reported here in full to document cross-species and cross-modality generalization of the toroidal geometry prediction.

#### Mouse hippocampal LFP (Figure 12)

The mouse hippocampal LFP provides a narrowband single-mode example. Its power spectral density contains a dominant theta-range oscillatory component superimposed on a broadband 1*/f* -like background (Figure 12c). Consistent with Proposition 2.1, a dominant sustained conjugate pole pair predicts *K* = 1 oscillatory mode and hence a one-dimensional toroidal skeleton T^1^ ≅ *S*^1^. Model-order selection via held-out MSE favors a low-order AR representation (Figure 12d), corresponding to one dominant oscillatory factor together with residual stable components that absorb broadband structure. The fitted model achieves strong held-out prediction accuracy (Figure 12b), confirming that the local AR approximation is appropriate within the analysis window. The recovered delay embedding organizes around a fitted one-torus (Figure 12f), indicating that the hippocampal LFP is well described by a single circular phase factor, with broadband fluctuations contributing geometric thickness rather than changing the intrinsic topology.

#### Mouse cortical EEG (Figure 13)

The mouse cortical EEG presents a qualitatively different regime. Its power spectral density is dominated by a broad 1*/f* -like aperiodic profile, with weaker and less isolated oscillatory components than the hippocampal LFP (Figure 13c). Nevertheless, model-order selection favors *p*^*⋆*^ = 4 (Figure 13d), consistent with two effective complex conjugate pole pairs and hence a two-mode local oscillatory representation. The recovered AR coefficients display time-varying oscillatory structure (Figure 13e), and the delay embedding is well described by a fitted two-torus (Figure 13f). This illustrates a key advantage of the delay-embedding perspective: multiple oscillatory components can be resolved as independent circular factors even when their spectral peaks are weak, overlapping, or not clearly separated in the PSD.

The cortical EEG also exhibits a larger Procrustes disparity than primate V1 LFP, reflecting weaker spectral separation and greater broadband contamination. This is consistent with the framework’s prediction that broadband or less sustained oscillatory activity produces thicker and less sharply resolved toroidal geometry. Together, the mouse hippocampal LFP and cortical EEG examples show that the proposed pipeline captures both single-mode and multi-mode oscillatory regimes across recording modalities, brain regions, and spectral profiles without dataset-specific tuning.

### E.2 Robustness Across Recording Channels in Monkey LFP

Figure 14a evaluates robustness across recording channels for primate V1 local field potential (LFP) recordings. For each channel, we compute the power spectral density (PSD), select the AR order using held-out prediction error, construct the lag embedding, and visualize the corresponding free-run AR prediction. Across channels, we consistently observe two dominant oscillatory components and an optimal AR order near *p*=4, corresponding to two complex conjugate pole pairs.

**Figure 12:**
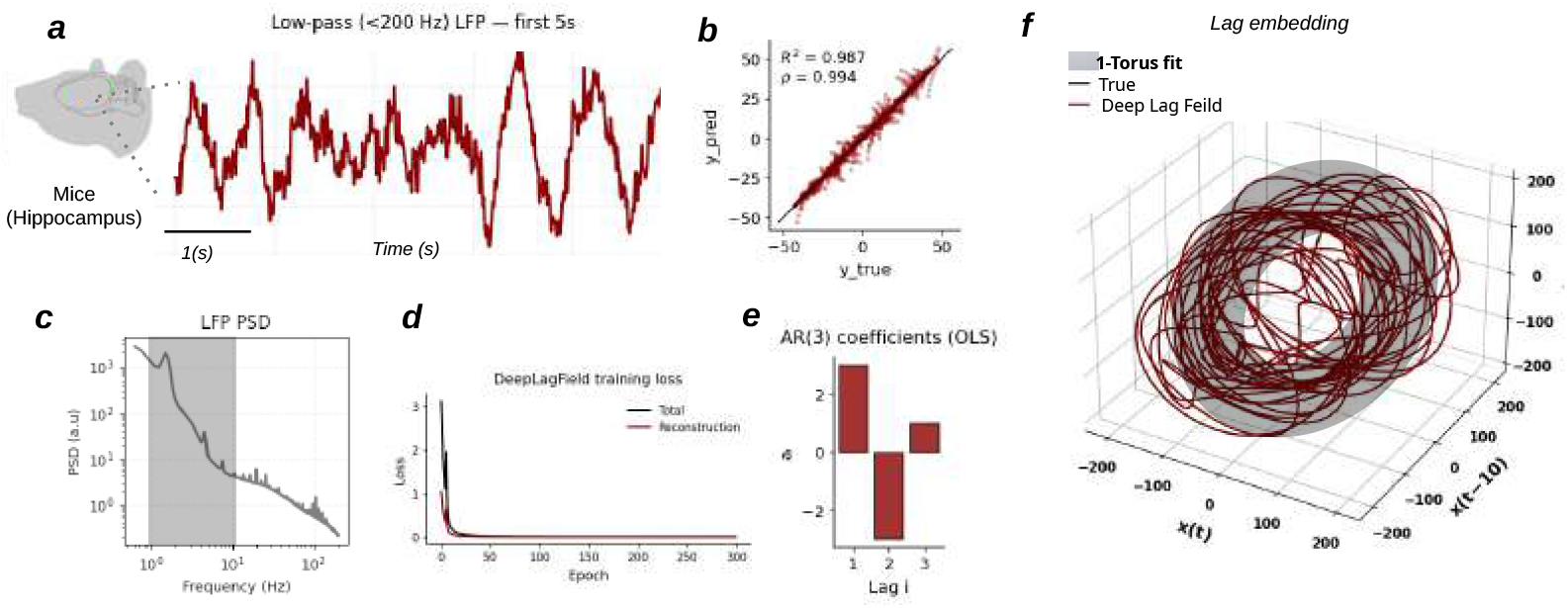
Manifold reconstruction of mouse hippocampal LFP. **(a)** Example low-pass filtered (*<* 200 Hz) LFP segment recorded from mouse hippocampus. **(b)** Held-out prediction performance, comparing predicted and true samples. **(c)** Power spectral density showing a dominant narrowband oscillatory component on top of a broadband 1*/f* -like background. **(d)** Held-out MSE as a function of AR order, selecting a low-order oscillatory model. **(e)** Estimated time-varying AR coefficients recovered by DeepLagField. **(f)** Delay embedding of the reconstructed hippocampal dynamics overlaid with a fitted 1-torus, consistent with a single dominant circular phase factor.

**Figure 13:**
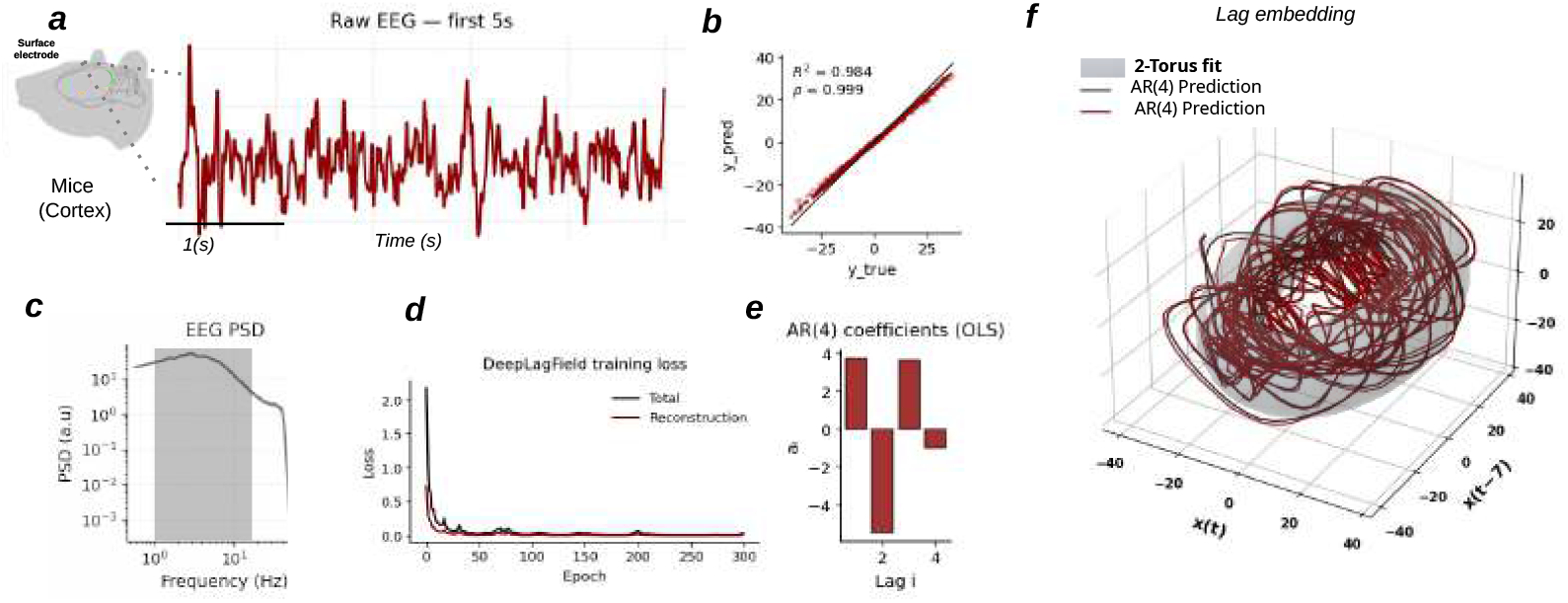
Manifold reconstruction of mouse cortical EEG. **(a)** Example EEG segment recorded from mouse cortex using a surface electrode. **(b)** Held-out prediction performance, comparing predicted and true samples. **(c)** Power spectral density showing a broadband 1*/f* -like profile with weaker narrowband structure than the hippocampal LFP. **(d)** Held-out MSE as a function of AR order, selecting *p*^*⋆*^ = 4. **(e)** Estimated time-varying AR coefficients recovered by DeepLagField. **(f)** Delay embedding of the reconstructed cortical EEG dynamics overlaid with a fitted 2-torus, consistent with two effective circular phase factors despite weak spectral peak separation.

**Figure 14:**
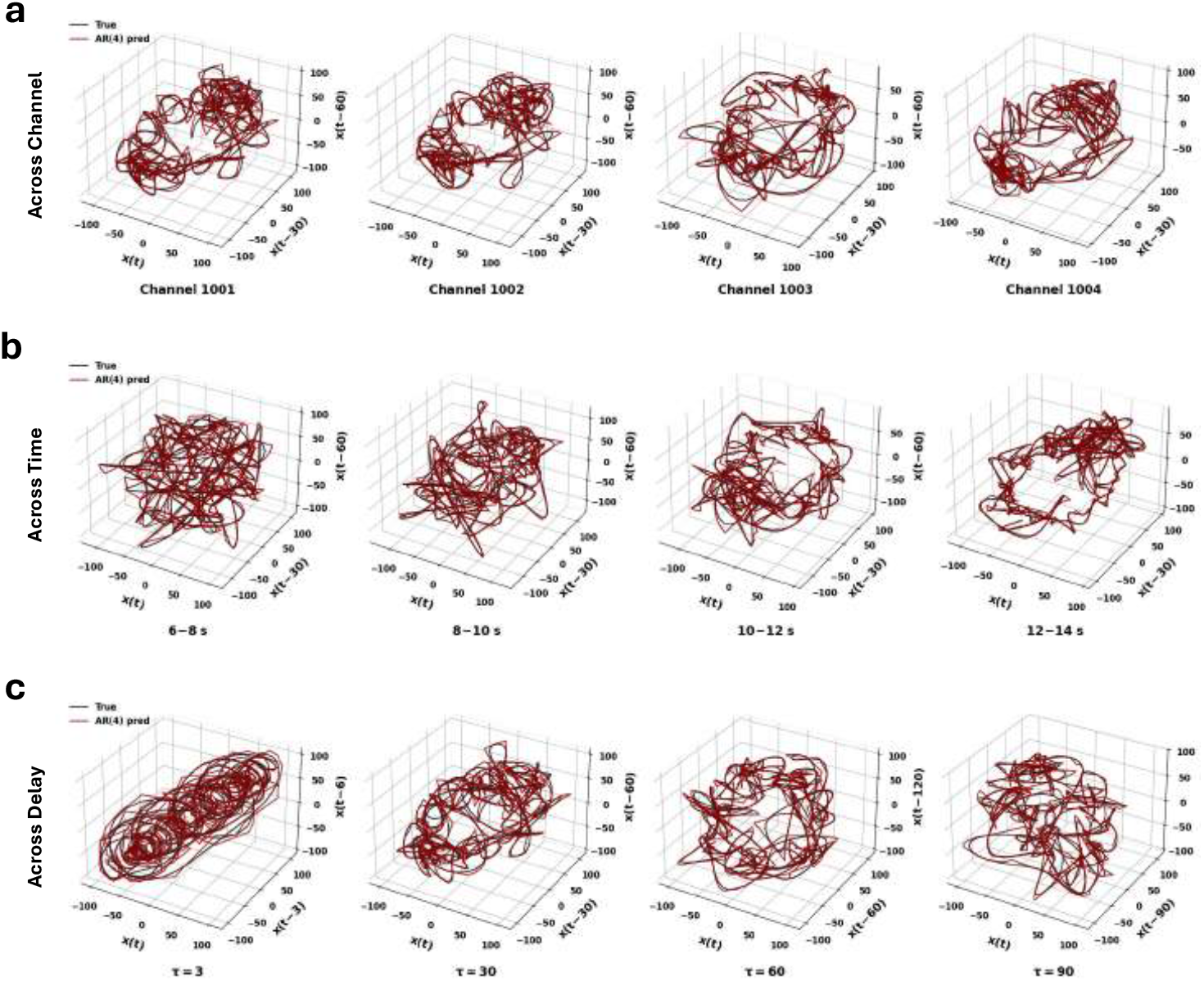
Robustness of toroidal geometry in monkey LFP recordings. **(a)** Lag embeddings and AR(4) free-run predictions for multiple channels recorded from primate V1. Across channels, the recovered geometry consistently exhibits a two–torus structure, supporting the interpretation that oscillatory content predicts embedding geometry at the population level. **(b)** Lag embeddings computed from different temporal windows of the same recording. Strong toroidal geometry appears in windows with sustained oscillatory structure and weakens when oscillations become transient or dominated by aperiodic background activity. **(c)** Lag embeddings constructed using different embedding delays *τ*. The two–torus geometry persists across a wide range of *τ*, degrading only for extremely small or large delays.

Despite variability in signal amplitude and noise characteristics across electrodes, the recovered lag embeddings exhibit qualitatively similar two-torus geometry. This consistency indicates that the inferred manifold structure reflects population-level dynamics rather than channel-specific artifacts.

### E.3 Local-in-Time Geometry in Nonstationary Monkey Recordings

Neural recordings are inherently nonstationary: oscillatory amplitudes and spectral content vary over time. To examine how geometry evolves locally, we apply the analysis pipeline to non-overlapping temporal windows within a single long monkey LFP recording.

Figure 14b shows that when two oscillatory components are simultaneously present within a window, the lag embedding exhibits a clear two-torus structure. In contrast, windows with weaker or transient oscillations produce degraded or partially collapsed geometry. These observations support the interpretation that toroidal structure is a local-in-time property governed by the instantaneous oscillatory content of the signal.

### E.4 Sensitivity to Delay Selection in Monkey LFP

The embedding delay *τ* determines the temporal separation between coordinates in the delay embedding. Figure 14c examines sensitivity to *τ* by sweeping a wide range of delay values.

While extreme choices of *τ* affect the visual compactness of the embedding, the qualitative two-torus geometry remains stable across a broad intermediate range. This robustness indicates that the geometric conclusions are not driven by fine-tuned delay selection.

### E.5 Cross-Species Validation: Mouse LFP Recordings

We repeat the same analyses on mouse LFP recordings collected independently of the primate dataset. In these recordings, spectral analysis typically reveals a single dominant oscillatory peak over the time windows considered. Accordingly, AR order selection frequently favors *p*=3, corresponding to a single complex conjugate pole pair.

Consistent with the theory, the resulting lag embeddings concentrate on disk-like structures rather than higher-dimensional tori. The analogous robustness analysis across channels, time windows, and embedding delays is shown in Figure 15: in each case the disk-like geometry is preserved wherever the single dominant theta oscillation is sustained, and degrades when that oscillation weakens, mirroring the pattern observed for monkey V1 but at a lower intrinsic dimensionality (*K*=1 vs. *K*=2). This parallel behavior across species provides additional confirmation that the relationship between oscillatory mode count and embedding topology is a modality-invariant property of the framework rather than a species- or region-specific artifact.

**Figure 15:**
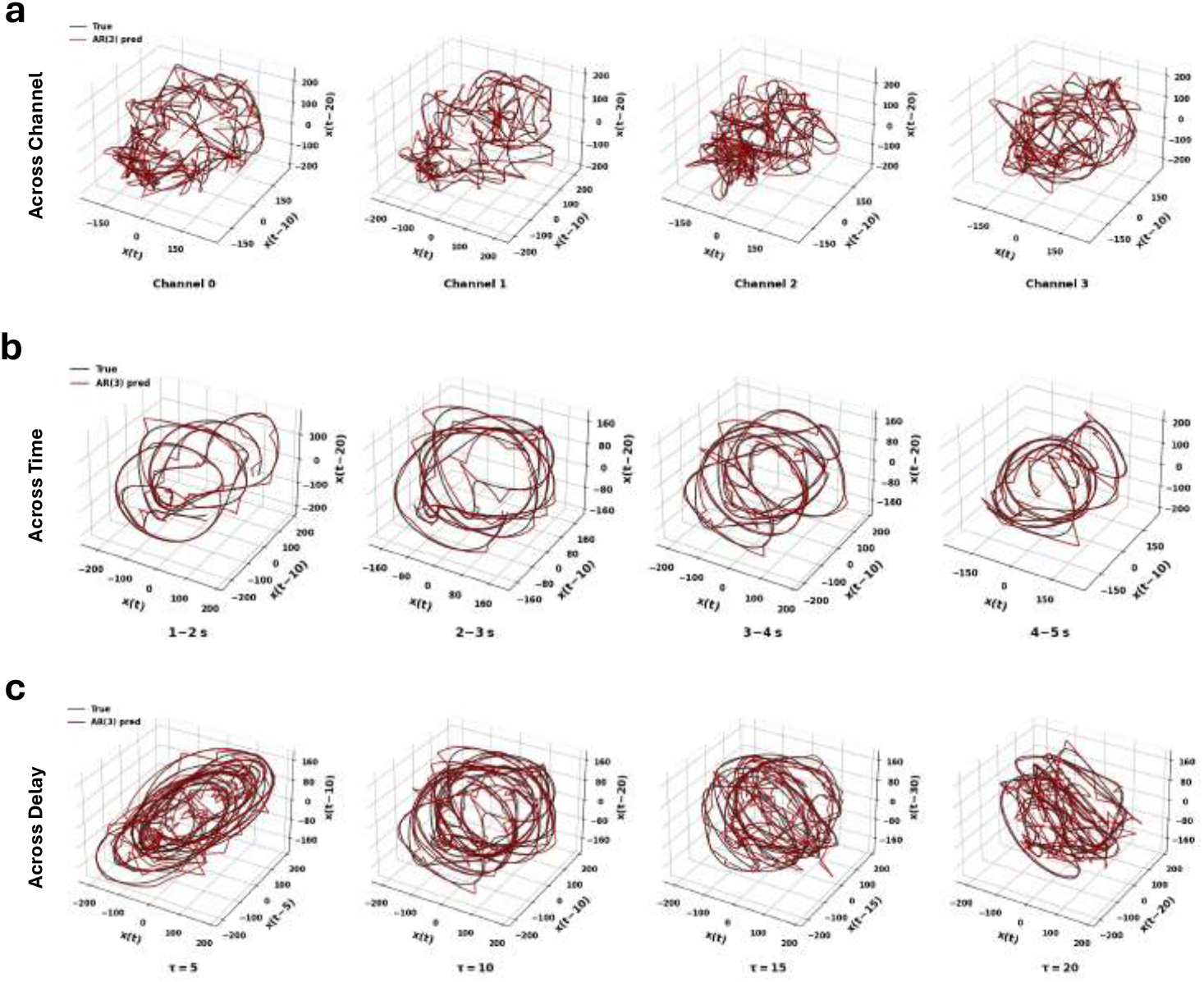
Mouse LFP recordings. Analogous analyses to the monkey results. Disk-like embeddings reflect the presence of a single dominant oscillatory mode.

## F Sensitivity and Robustness Analysis

This appendix evaluates the robustness of the recovered lag-embedded geometry to key analysis choices, including delay *τ*, window length *T*_*w*_, and AR order selection. The goal is to verify that the qualitative conclusions of the paper are stable across reasonable hyperparameter choices and that observed failure modes align with theoretical expectations rather than tuning artifacts.

### F.1 Sensitivity to Window Length *T*_*w*_

The window length *T*_*w*_ governs a bias–variance tradeoff between local stationarity and estimation accuracy. Across a wide range of window lengths, the selected AR order and inferred embedding geometry remain stable. Extremely short windows increase estimation noise, while very long windows may mix distinct dynamical regimes, but intermediate values yield consistent results.

### F.2 Stability of AR Order Selection

Across windows and recordings, AR order selection concentrates tightly around values predicted by the number of dominant oscillatory modes. Monkey recordings favor *p*≈ 4, while mouse recordings favor *p*≈ 2. This stability reinforces the interpretation that AR order captures the number of active phase variables rather than incidental signal fluctuations.

#### Summary

Across species, channels, and time windows, the recovered lag-embedded geometry is robust to reasonable choices of delay and window length. Toroidal or disk-like structures emerge precisely when sustained oscillatory components are present and degrade when they are absent or transient. These results support the central claim that oscillatory structure predicts the local geometry of delay embeddings in real neural recordings.

## G Datasets

We evaluate the proposed framework on a combination of biological and synthetic datasets designed to assess both theoretical predictions and practical robustness under realistic recording conditions. The biological datasets provide empirical validation across species and acquisition modalities, while the synthetic datasets enable controlled evaluation under known ground-truth dynamics. Together, these datasets allow systematic assessment of topology preservation, reconstruction accuracy, and robustness to nonstationarity.

### G.1 Biological datasets

We consider three electrophysiological datasets spanning invasive and non-invasive neural recordings: invasive local field potential (LFP) recordings from non-human primates, invasive LFP recordings from mice, and chronic electroencephalography (EEG) recordings from mice. Across all biological datasets, our analysis operates on single-channel time series in order to isolate intrinsic temporal geometry and avoid confounds arising from spatial mixing or multichannel aggregation.

#### Non-human primate LFP recordings

We used a publicly available macaque visual cortex electrophysiology dataset comprising resting-state recordings from rhesus macaques (*Macaca mulatta*) implanted chronically with a 1024-channel intracortical interface spanning V1 and V4 Chen et al. [2022]. The full implant consisted of 16 Utah microelectrode arrays (64 electrodes per array; 8× 8 geometry), with dense coverage in V1 and additional coverage in V4 Chen et al. [2022]. Signals include broadband extracellular recordings (sampled at 30 kHz) and derived local field potential (LFP; 1–100 Hz) representations, with simultaneous eye tracking during head-fixed sessions Chen et al. [2020].

In this study, we restricted analyses to recordings from a *single V1 Utah array* (one 64-electrode array) to standardize spatial scale and to focus on single-electrode temporal geometry. All analyses were performed on single-electrode LFP time series (no multichannel aggregation). We applied a consistent preprocessing pipeline (band-limiting to the LFP range, removal of line-noise components when present, and resampling to a common rate) prior to delay-embedding and downstream evaluation.

#### Mouse LFP recordings

A four-channel linear silicon probe (Q-trode, NeuroNexus) was chronically implanted to record hippocampal LFP, contralateral to the imaging window. Surgical procedures largely followed those described previously Grosmark et al. [2021]. Briefly, a 0.7-mm burr hole was drilled at coordinates 2.2 mm AP) and 2.75 mm ML relative to bregma. The probe was lowered to a depth of 1.0 mm DV relative to the cortical surface. The probe was inserted at a 45° angle and secured to the skull with dental cement for chronic recordings. A reference screw was placed over the cerebellum. After post-surgical recovery (*>*5 days), the mice were habituated to the head-fixed tread-mill setup (5–7 days). Simultaneous LFP recordings and in vivo calcium imaging were performed; however, only the LFP data were analyzed in this study. Signals were acquired at 20 kHz using a multiplexed digital acquisition system (Intan RHD).

#### Mouse EEG recordings

Mice were implanted for chronic EEG and EMG recordings under ketamine/xylazine anesthesia (100 and 10 mg ·kg^*−*1^, intraperitoneal) using stereotaxic surgery [Turi et al., 2025]. For EEG acquisition, stainless steel skull screws were positioned over the cortex approximately 1 mm lateral to the midline, with one screw placed 1.5 mm anterior to bregma and a second placed 1.5 mm posterior to bregma. A reference screw was positioned over the cerebellum.

For EMG recordings, two electrodes were inserted bilaterally into the neck musculature. EEG screws and EMG electrodes were connected to a printed circuit board (PCB) soldered to a five-position pin connector and secured to the skull using dental cement. Following surgery, animals were allowed to recover for a minimum of two weeks prior to data collection.

Sleep–wake behavior was monitored continuously for 24 hours in a sound-attenuating chamber following at least four hours of habituation. Behavioral monitoring was performed using infrared video recording at 30 frames per second, with a 12-hour light/dark cycle (lights on at 7:00 AM, off at 7:00 PM). EEG and EMG signals were bandpass filtered between 0.5 and 500 Hz and digitized at 1017 Hz using multichannel amplifiers (TDT PZ5 and RZ5D or Neuralynx Digital Lynx 4S).

### G.2 Synthetic datasets

In addition to biological recordings, we evaluate the proposed method on synthetic time series generated from controlled autoregressive (AR) models. These datasets are used to isolate specific dynamical properties, evaluate identifiability under known ground truth, and assess robustness to nonstationarity in a setting where the intrinsic manifold geometry is analytically tractable.

#### Stationary AR processes

We generate synthetic signals from fixed-coefficient autoregressive models of finite order,

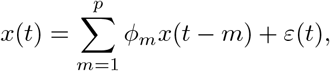

where the coefficients {*ϕ*_*m*_}are constant in time and chosen to produce stable dynamics with pre-scribed oscillatory content. Innovation noise *ε*(*t*) is zero-mean and temporally white. For each stationary AR realization, 100,000 samples are generated, providing sufficient data to accurately estimate spectral structure and delay-embedded geometry. These datasets directly match the assumptions underlying the analytical results in Section 3 and serve as a controlled sanity check.

#### Time-varying AR processes

To evaluate performance under nonstationarity, we generate time-varying autoregressive (TVAR) processes of the form

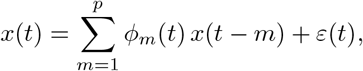

where the coefficients {*ϕ*_*m*_(*t*)} evolve smoothly over time. For each realization, 100,000 samples are generated. Coefficient trajectories are drawn from a diverse family of smooth function classes designed to capture different forms of temporal variability observed in neural data.

#### Classes of time-varying coefficient trajectories

To systematically evaluate reconstruction accuracy, order stability, and geometric robustness under different forms of nonstationarity, we generate TVAR benchmarks using a diverse set of smooth coefficient trajectories. Specifically, we consider the following.

1. sinusoidal modulations,
2. finite Fourier series with multiple harmonics,
3. quasi-periodic combinations of incommensurate frequencies,
4. low-order polynomial drifts,
5. logistic transitions modeling regime switches,
6. superpositions of smooth sigmoid or Gaussian-shaped bumps,
7. and smooth stochastic trajectories obtained by filtering random processes.

All coefficient schedules are constructed to ensure stability of the resulting AR dynamics at every time step (i.e., all instantaneous poles remain inside the unit disk). This controlled design allows us to isolate the effect of different nonstationarity patterns on coefficient recovery, effective order selection, and preservation of the underlying delay-embedding geometry.

Representative examples of coefficient trajectories from each function class are shown in Figure 16, illustrating the range of temporal behaviors considered.

**Figure 16:**
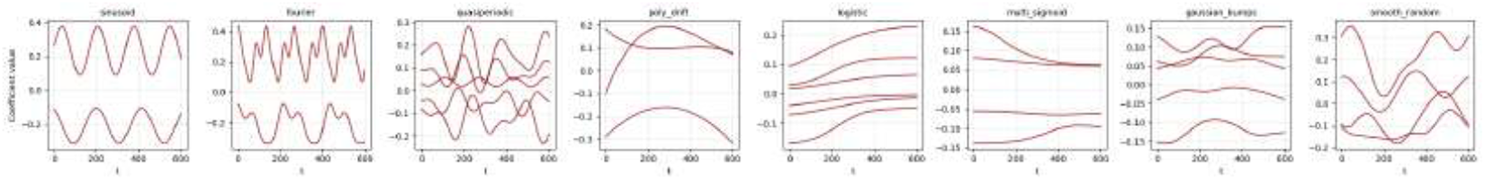
Representative time-varying autoregressive coefficient trajectories used to generate synthetic TVAR benchmarks. Each panel corresponds to a different class of nonstationarity, including sinusoidal, Fourier, quasi-periodic, polynomial drift, logistic transitions, multi-sigmoid, Gaussian bump, and smooth stochastic schedules. All trajectories are constructed to preserve stability of the induced AR dynamics.

#### Common preprocessing and evaluation protocol

To ensure comparability across biological and synthetic datasets, all signals are processed using a consistent preprocessing and embedding pipeline. Signals are filtered, downsampled to a common sampling rate, segmented into locally stationary windows, and normalized to remove amplitude scaling effects. No behavioral labels or task variables are used during model fitting; all analyses operate directly on the neural time series. This unified protocol allows direct comparison of geometric recovery across controlled synthetic systems and real neural recordings.

## H Additional Ablation Results

This appendix reports full ablation results complementing the summary presented in the main text. All experiments are conducted on the synthetic time-varying autoregressive (TVAR) benchmarks described in Appendix G. Each ablation removes a single component from the full DeepLagField model while keeping the remaining architecture, training data, and optimization procedure fixed. Performance is evaluated using TVAR coefficient mean-squared error (MSE), one-step signal prediction MSE, and autoregressive order recovery accuracy.

Across ablations, distinct failure modes emerge. Removing the autoregressive structure results in large increases in both coefficient and signal prediction error. Removing temporal smoothness substantially degrades coefficient estimation despite only modest changes in prediction error, indicating that smoothness primarily stabilizes nonstationary identification. Most notably, removing the order-consistency constraint yields near-identical prediction error to the full model, yet causes a collapse in order recovery accuracy across most orders. This confirms that predictive accuracy alone is insufficient for identifying intrinsic temporal structure and motivates the geometric constraints imposed by the full model.

## I Training Procedure for DeepLagField

This appendix details the training objective and optimization procedure used for *DeepLagField*. Both models are trained end-to-end to infer a time-varying autoregressive (TVAR) representation of neural time series while jointly selecting the effective AR order. The distinction between the two models lies solely in the loss function: *DeepLagField* augments the reconstruction objective with physics-informed regularization terms, whereas *DeepTori* uses only the prediction loss.

### I.1 Model inputs and outputs

Given a minibatch of *N* time series segments

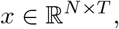

the network outputs: (i) a discrete estimate of the effective AR order 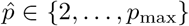, (ii) time-varying AR coefficients 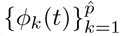, and (iii) a one-step-ahead reconstruction 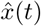. Discrete order selection is implemented using a Gumbel–Softmax relaxation, allowing gradient-based optimization with a straight-through estimator.

### I.2 Training objective

#### Reconstruction loss

Let 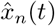 denote the reconstructed signal for sequence *n* at time *t*, obtained from the predicted coefficients via

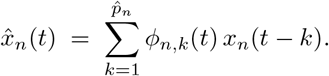

The base reconstruction loss is

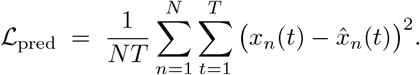

#### Smoothness regularization (local stationarity)

To encode the assumption that neural dynamics evolve gradually over short time intervals, we penalize temporal variation in the AR coefficients:

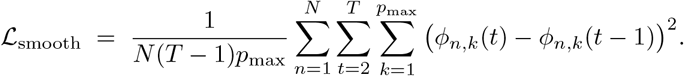

#### Bounded-energy regularization

LFP signals empirically exhibit bounded amplitude and smoothly varying power over short temporal windows. We define a local energy proxy

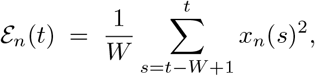

where *W* is a fixed window length. We penalize rapid energy fluctuations via

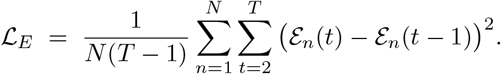

#### Total loss

The full training objective for *DeepLagField* is

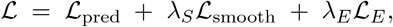

where *λ*_*S*_, *λ*_*E*_ ≥ 0 are fixed hyperparameters. For *DeepTori*, we set *λ*_*S*_ = *λ*_*E*_ = 0.

### I.3 Optimization

All parameters are trained using stochastic gradient descent with the Adam optimizer. Gradients are propagated through the discrete order-selection mechanism via the straight-through estimator induced by the Gumbel–Softmax relaxation.

### I.4 Training algorithm

Algorithm 1 summarizes the training procedure for one minibatch.

#### Algorithm 1

**DeepLagField training per minibatch**

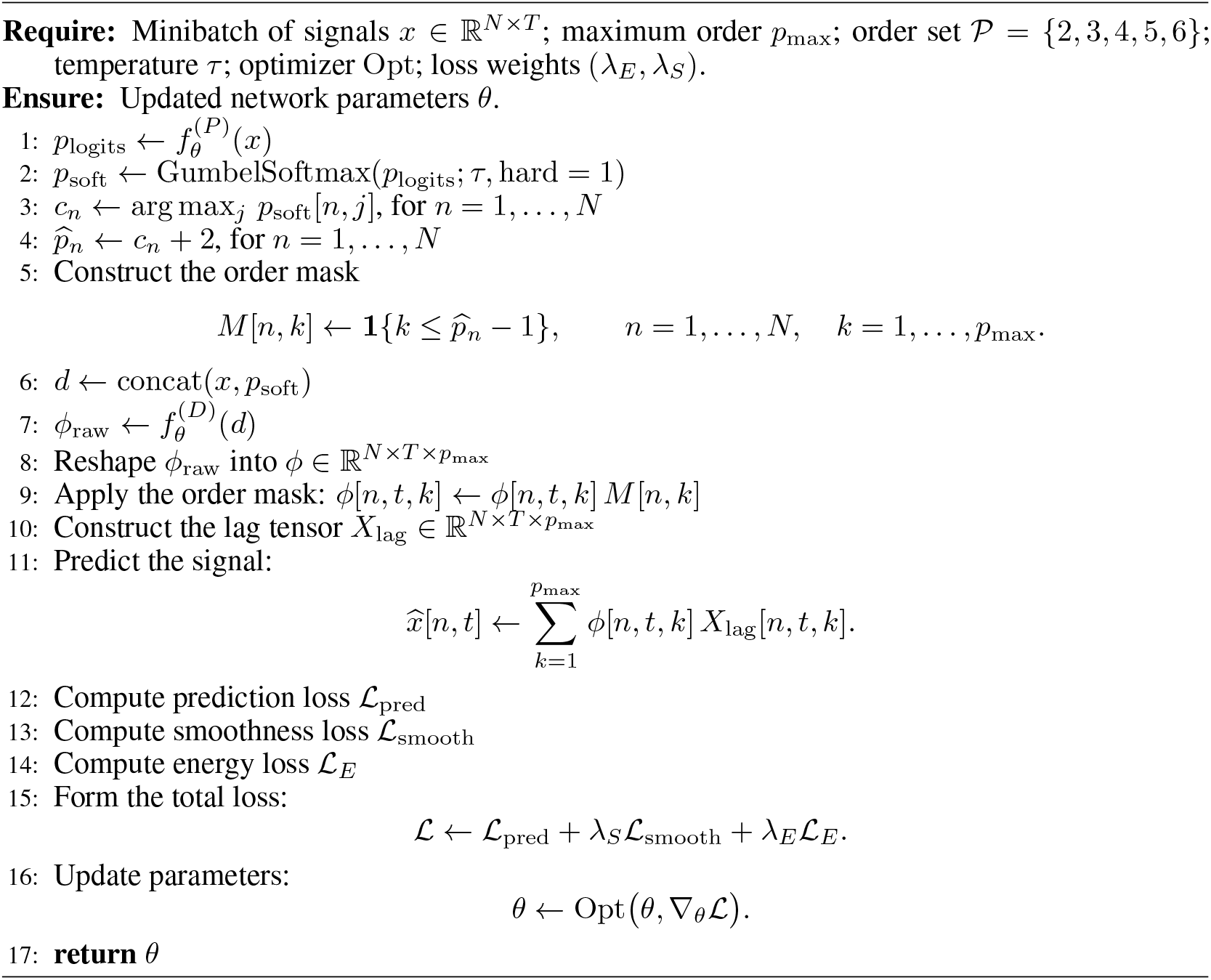

## J Geometry of Lag Embeddings Under Aperiodic 1*/f* ^*α*^ Noise

This appendix provides additional intuition and supporting simulations clarifying the geometric role of aperiodic 1*/f*^*α*^ structure in lag–embedded time series. These results complement the main analysis by illustrating how aperiodic components influence the *conditioning* and *thickness* of embeddings, while leaving intrinsic topology unchanged. As emphasized in the main text, oscillatory components determine the number of circular factors in the manifold, whereas aperiodic structure modulates geometric regularity and robustness.

### J.1 Spectral and Temporal Properties of 1*/f*^*α*^ Processes

We consider zero–mean stationary Gaussian processes whose power spectral density (PSD) scales as

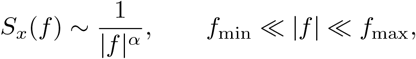

with *α* ∈ {0, 1, 2, 3}. Figure 17a illustrates the associated PSDs, autocorrelation functions, and representative time series realizations.

**Figure 17:**
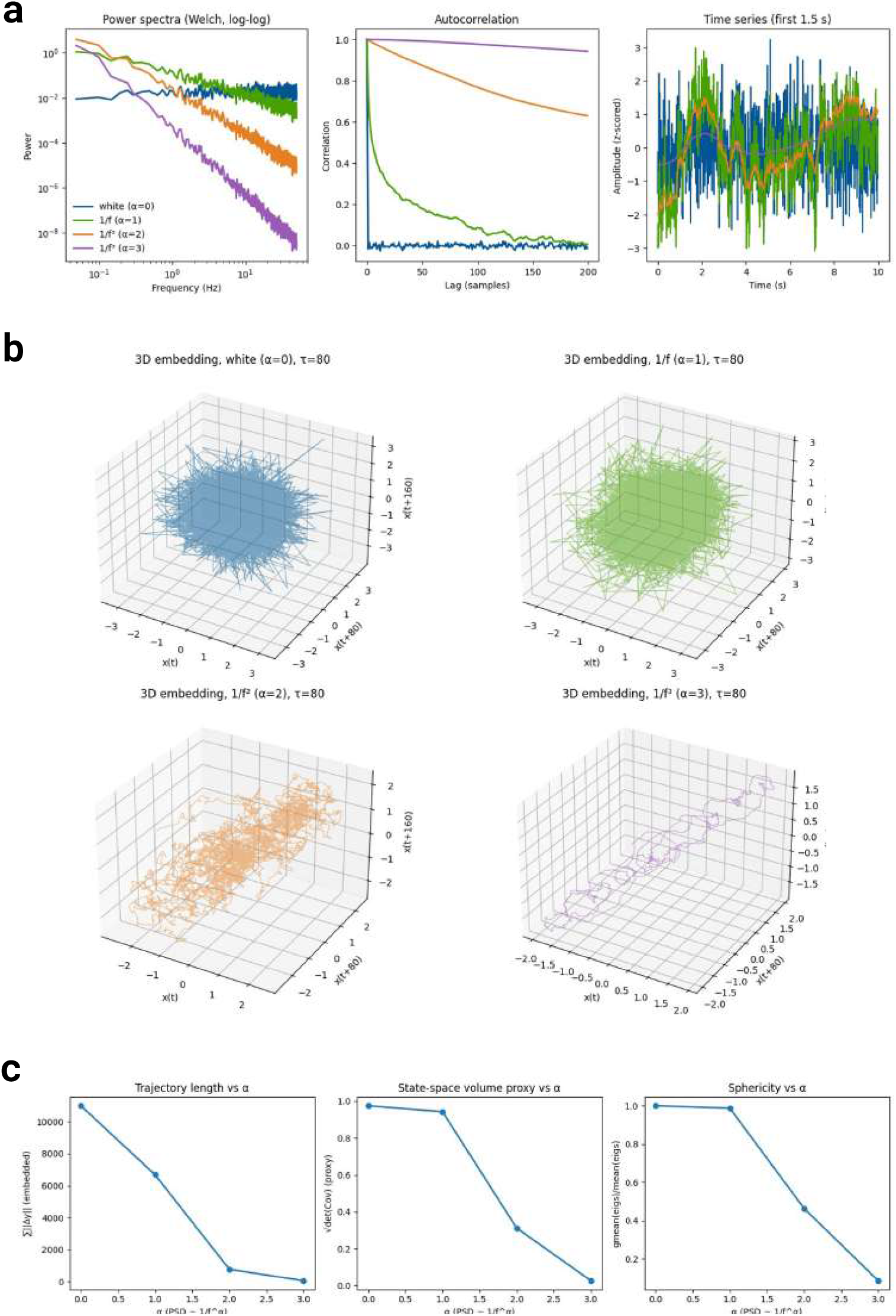
Spectral, temporal, and geometric structure of 1*/f*^*α*^ noise. **(a)** Power spectral densities (Welch estimate, log–log scale), autocorrelation functions, and representative time series realizations (z–scored) for *α* = 0, 1, 2, 3. Increasing *α* induces long–range temporal correlations and smoother trajectories. **(b)** Three–dimensional lag embeddings (*τ* = 80). White noise (*α* = 0) yields an isotropic cloud; 1*/f* noise (*α* = 1) produces correlated but space–filling geometry; 1*/f* ^2^ noise collapses toward a low–dimensional filament; 1*/f* ^3^ noise exhibits near–curve–like trajectories. Increasing *α* drives geometric degeneration without creating circular structure. **(c)** Quantitative geometric measures as a function of *α*: embedded trajectory length (left), state–space volume proxy 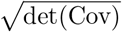 (middle), and sphericity (geometric mean / arithmetic mean of covariance eigenvalues; right). All three measures decrease monotonically with *α*, confirming systematic geometric collapse and growing anisotropy.

**Figure 18:**
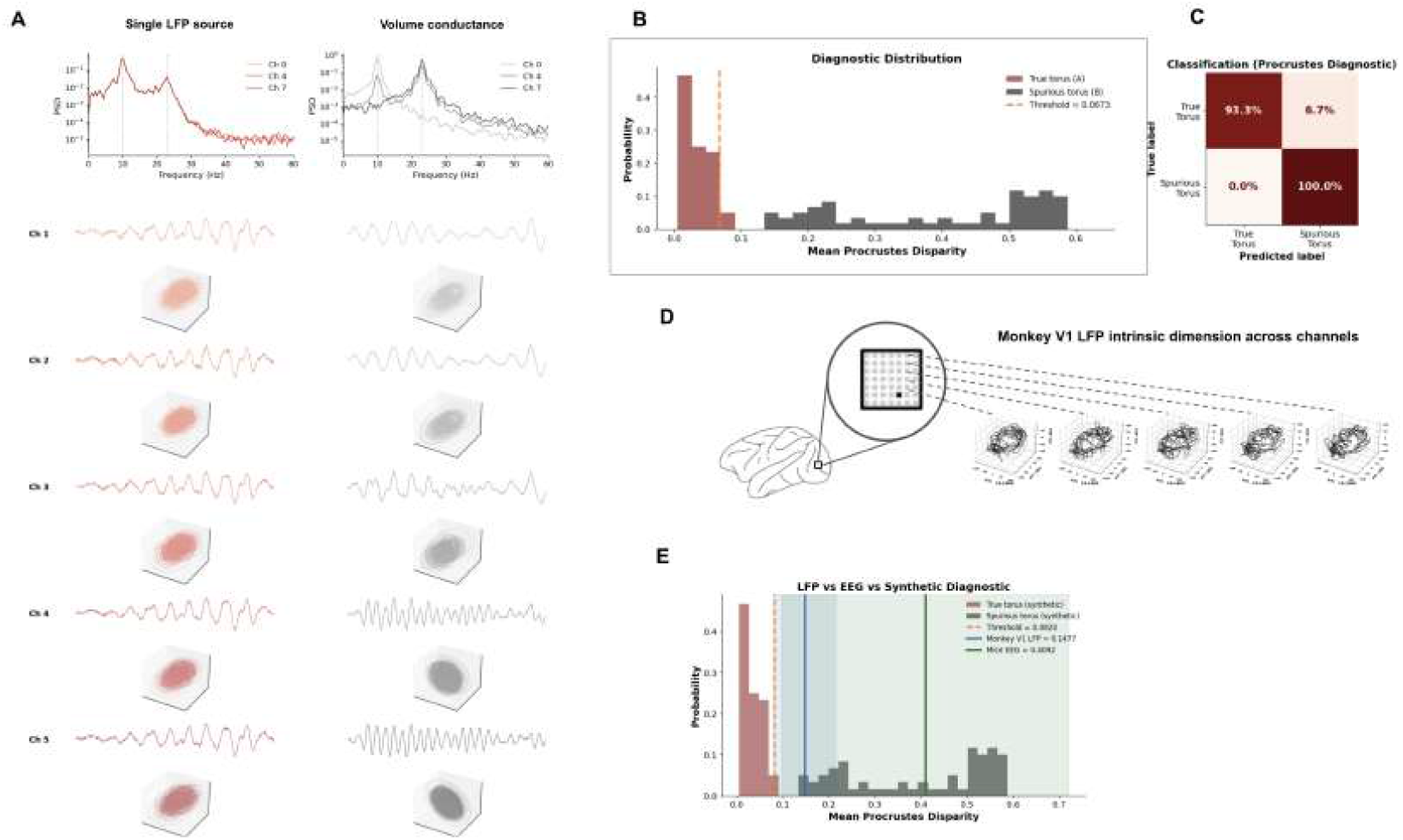
Multichannel analysis and Procrustes disparity diagnostic. **(A)** Comparison of single LFP source (left) versus volume conduction (right) conditions. Top: power spectral densities for representative channels (Ch 0, Ch 4, Ch 7) showing similar spectral profiles across conditions. Bottom: lag embeddings recovered from five individual channels (Ch 1–5) under each condition. Single source recordings yield clean toroidal or disk-like embeddings (red), while volume-conducted signals produce degraded ellipsoidal geometry (gray), confirming that toroidal structure reflects genuine local oscillatory dynamics. **(B)** Distributions of mean Procrustes disparity for synthetic true torus signals (dark red) and spurious torus signals (gray). The two distributions are well-separated, with a diagnostic threshold of 0.0673 (orange dashed line) providing near-optimal classification. **(C)** Confusion matrix for the Procrustes disparity classifier on held-out synthetic signals. The classifier achieves 93.3% accuracy on true torus signals and 100% accuracy on spurious torus signals. **(D)** Cross-channel validation in monkey V1. Lag embeddings recovered from four representative channels of the Utah array (Ch 1001–1004) show consistent two-torus geometry across channels, confirming that the recovered manifold structure reflects population-level dynamics rather than channel-specific artifacts. **(E)** Procrustes disparity diagnostic applied to real recordings, overlaid on synthetic calibration distributions. Monkey V1 LFP (blue line; disparity = 0.1477) falls above the true torus threshold (0.0820, orange dashed) but below the spurious torus bulk, indicating valid toroidal geometry with moderate noise. Mouse cortex EEG (green line; disparity = 0.4092) falls within the spurious torus distribution, consistent with its noisier spectral structure and the absence of a dominant narrowband oscillatory peak. The diagnostic correctly grades geometric quality on a continuous scale rather than producing binary classifications.

For *α* = 0 (white noise), samples are temporally uncorrelated and the autocorrelation decays immediately. As *α* increases, the PSD becomes increasingly dominated by low frequencies, leading to progressively longer temporal correlations. In the time domain, this manifests as smoother trajectories with pronounced long–range memory. These effects are well known from the theory of long–memory stochastic processes, but their geometric consequences for delay embeddings are often left implicit. For a complementary information-theoretic treatment of temporally correlated Gaussian noise, particularly Gauss–Markov processes, see Fallah [2024].

### J.2 Lag–Embedded Geometry

To visualize the geometric effect of aperiodic structure, we construct fixed–lag embeddings

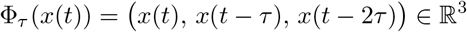

with lag *τ* chosen beyond the short–term correlation scale. Figure 17b shows three–dimensional embeddings for different values of *α*.

White noise (*α* = 0) produces an approximately isotropic point cloud filling a ball in state space. As *α* increases, correlations between delayed coordinates induce strong anisotropy: the embedding collapses toward elongated, filamentary, or nearly one–dimensional structures. For *α* = 2 and *α* = 3, the trajectory closely resembles integrated noise, exhibiting smooth, drift–dominated paths rather than space–filling clouds.

Importantly, this geometric collapse occurs *without introducing new topological features*. Unlike oscillatory components, which generate circular phase variables and hence toroidal topology, aperiodic 1*/f*^*α*^ structure produces no closed loops or persistent homology in degree one. Instead, it controls the effective dimensionality and conditioning of the embedding.

### J.3 Quantitative Geometric Degeneration

To quantify this transition, we compute three geometric proxies from the embedded trajectories: (i) total trajectory length, (ii) a covariance–based volume proxy 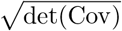, and (iii) sphericity, defined as the ratio between the geometric and arithmetic means of the covariance eigenvalues. Results are summarized in Figure 17c.

All three measures decrease monotonically with *α*, confirming that increasing spectral exponent leads to progressively more degenerate embeddings. White noise yields maximal volume and near–unit sphericity, while large *α* produces elongated, low–volume structures. These effects directly impact the empirical recoverability of manifold geometry in noisy data.

### J.4 Interpretation and Relation to the Main Results

These simulations clarify the distinct roles of oscillatory and aperiodic components in lag–embedded geometry. Oscillatory peaks correspond to near–unit–modulus conjugate poles and generate circular phase variables, determining the intrinsic topology of the manifold. In contrast, aperiodic 1*/f*^*α*^ structure introduces long–range temporal correlations that affect smoothness, anisotropy, and noise sensitivity, but do not contribute additional topological dimensions.

This separation explains why spectral equalization can substantially improve the empirical visibility of toroidal manifolds without altering Betti numbers: equalization rescales anisotropic directions induced by amplitude imbalance and aperiodic structure, while preserving the underlying product–of– circles topology dictated by oscillatory modes.

## K Compute, Implementation, and Scalability Details

### K.1 Compute and Implementation

#### Implementation Details

All experiments were implemented in PyTorch using standard autograd for gradient computation. Training was performed on a workstation equipped with dual NVIDIA GeForce RTX 5090 GPUs (32 GB VRAM; Blackwell architecture), an AMD Ryzen Threadripper PRO 9975WX processor (64 cores @ 5.484 GHz), and 512 GB system RAM, running Ubuntu 24.04.3 LTS with Linux kernel 6.8.0. The DeepLagField model with default configuration (*d*=128, *p*_max_=6) contains 652,565 trainable parameters.

The synthetic benchmark dataset comprised 100,000 time-varying autoregressive (TVAR) samples spanning eight function families: sinusoidal modulation, finite Fourier series, quasi-periodic functions, low-order polynomial drift, logistic transitions, Gaussian-shaped bumps, piecewise-linear ramps, and smooth stochastic trajectories. Empirical validation was performed on primate V1 local field potentials (LFP), rodent hippocampal LFP, and mouse EEG recordings. DeepLagField was trained for 200 epochs with a batch size of 32 using the Adam optimizer with learning rate 10^*−*3^.

A single optimization step required approximately 0.83 ms on a single RTX 5090 GPU, with inference latency of 0.30 ms per batch. Total training time scaled linearly with the number of batches per epoch and the number of epochs. Training remained stable throughout all experiments, with no observed gradient explosions or numerical instabilities.

### K.2 Scalability with Respect to Network Width

We evaluated how training behavior and computational cost scale with hidden dimension *d* as a controlled robustness check. This experiment is not intended as a competitive efficiency benchmark, but rather as a sanity check demonstrating that the method does not rely on narrow architectural regimes.

Increasing the hidden dimension by 16 × (from 32 to 512) resulted in an ∼19× increase in trainable parameters (159,701 to 2,992,661), yet training time per step remained nearly constant at 0.80–0.85 ms and inference latency at 0.30–0.32 ms. This behavior confirms that the dominant computational cost arises from sequence-length operations rather than network width, and that the architecture scales gracefully within the evaluated regime without sacrificing training stability. We note that this favorable scaling is specific to the current parameter range; asymptotic complexity will dominate at substantially larger model sizes.

## L Multichannel Analysis and Procrustes Diagnostic

This appendix provides full details of two complementary analyses supporting the main text: (i) the effect of volume conduction on recovered lag-embedded geometry in multichannel LFP recordings, and (ii) the Procrustes disparity diagnostic used to distinguish genuine toroidal geometry from spurious torus fits. Together these results establish that the toroidal manifold structure recovered in the main text reflects genuine oscillatory dynamics rather than spatial mixing artifacts or fitting coincidences, and provide a principled tool for grading geometric quality in new recordings.

### L.1 Volume Conduction and Single-Channel Geometry

A fundamental concern in LFP analysis is that signals recorded across nearby electrodes may reflect the same underlying source through volume conduction rather than independent local dynamics. This subject is more vital when studying the collective dynamics of neural field in multichannel domains and relevant wave-based analysis Das et al. [2023, 2026]. If volume conduction dominates, the geometry recovered from individual channels may be artifactual — reflecting spatial mixing rather than intrinsic oscillatory structure. We examine this directly by comparing lag embeddings from simulated single-source LFP signals against signals corrupted by volume conductance.

#### Simulation setup

We simulate two conditions. In the *single source* condition, each channel receives an independent draw from the same AR process, preserving the oscillatory structure without spatial mixing. In the *volume conduction* condition, channels receive a weighted mixture of a common source with distance-dependent attenuation, modeling the spatial smearing characteristic of extracellular recordings. Power spectral densities and raw time series for representative channels (Ch 0, Ch 4, Ch 7) are shown in Figure 18A for both conditions.

#### Geometric consequences

Figure 18A shows the lag embeddings recovered from individual channels under both conditions across five representative channels (Ch 1–5). Under the single source condition (left column), all channels recover clean toroidal or disk-like embeddings consistent with the predicted oscillatory manifold structure. Under volume conduction (right column), the embeddings collapse toward ellipsoidal or disk-like structures with degraded toroidal geometry, reflecting the loss of independent phase information across channels due to spatial mixing.

These results have two important implications. First, they confirm that clean toroidal geometry in single-channel recordings is a signature of genuine local oscillatory dynamics rather than a trivial consequence of spatial structure. Second, they show that volume conduction degrades but does not destroy geometric structure — the embeddings remain low-dimensional and bounded, consistent with the theoretical prediction that aperiodic and mixed components contribute thickness rather than new topological dimensions (Section 2).

### L.2 Procrustes Disparity Diagnostic

A key practical challenge in applying our framework to real data is distinguishing genuine toroidal geometry from spurious torus fits that arise from noise or aperiodic structure. We address this using a Procrustes disparity diagnostic: a data-driven threshold calibrated on synthetic signals with known ground-truth geometry.

#### Diagnostic construction

We generate two classes of synthetic signals. *True torus* signals (class A) are drawn from AR processes with sustained oscillatory poles, producing genuine toroidal lag embeddings. *Spurious torus* signals (class B) are drawn from AR processes with only stable aperiodic poles, producing embeddings that may superficially resemble tori under finite-sample noise but lack genuine circular topology. For each signal, we fit a torus to the lag embedding and compute the mean Procrustes disparity between the fitted torus and the point cloud — a measure of how well the torus model accounts for the observed geometry.

#### Threshold calibration

Figure 18B shows the resulting distributions of mean Procrustes disparity for true torus (dark red) and spurious torus (gray) signals. The two distributions are well-separated, with true torus signals concentrated near zero disparity and spurious torus signals exhibiting large, diffuse disparity values. A threshold of 0.0673 provides near-optimal separation between the two classes.

#### Classification performance

Figure 18C shows the confusion matrix for the Procrustes diagnostic classifier applied to held-out synthetic signals. The classifier achieves 93.3% accuracy on true torus signals and 100% accuracy on spurious torus signals, confirming that mean Procrustes disparity is a reliable and practically useful criterion for distinguishing genuine from spurious toroidal geometry.

### L.3 Cross-Channel Validation in Monkey V1

Figure 18D shows lag embeddings recovered from four representative channels of the monkey V1 Utah array. Consistent with the single-source simulation results in Appendix L.1, all channels recover qualitatively similar toroidal geometry, confirming that the manifold structure reflects population-level oscillatory dynamics rather than channel-specific artifacts. The intrinsic dimension estimated from each channel is consistent with the two-torus prediction of Section 2, with optimal AR order concentrating near *p* = 4 across channels.

## M Quantification of Toroidal Geometry

To assess the quality of recovered toroidal geometry, we employ two complementary quantification approaches: a geometric approach based on *torus score* and *torus error*, which measure how well the lag-embedded trajectory is captured by a fitted torus model, and a topological approach based on *windowed persistent homology*, which provides a coordinate-free verification of the underlying homotopy type. Together, these measures provide both a continuous summary of geometric fit quality and a discrete topological invariant that can be compared against the theoretical Betti-number predictions of Section 2.

### M.1 Torus Score and Torus Error

For each recording window, let 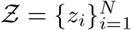 denote the lag-embedded trajectory, with *z*_*i*_ ∈ ℝ^*q*^. We fit the toroidal geometry predicted by the recovered AR dynamics and denote its fitted geometric region by 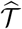. This region is allowed to have finite thickness, since empirical trajectories are not expected to lie exactly on an ideal torus in the presence of noise, broadband fluctuations, and finite-sample variability.

For each embedded point, we define a nonnegative residual

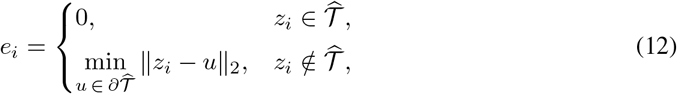

where 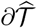 denotes the boundary of the fitted torus region. Points already contained within the recovered toroidal geometry contribute zero residual, while points outside contribute their shortest distance back to the torus boundary. The *torus score* is the fraction of embedded points contained by the fitted torus,

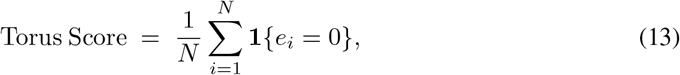

and the *torus error* is the mean residual distance,

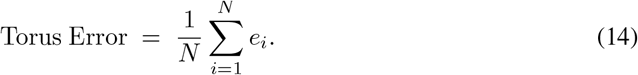

Intuitively, torus score measures how much of the lag-embedded trajectory is captured by the recovered toroidal geometry, whereas torus error measures the average distance by which the remaining points fall outside that geometry. A high torus score together with a low torus error indicates that the trajectory is well explained by the fitted torus. Torus error is reported in the units of the embedding coordinates and is interpreted under the same preprocessing and normalization used for the corresponding embedding.

#### Torus fitting procedure

The torus fit is initialized from the two leading principal components of the embedded point cloud, which provide an initial estimate of the major and minor radii (*R*_1_, *R*_2_) and the torus center. The parameters are then refined by minimizing the mean squared distance from the point cloud to the torus surface via gradient descent. To prevent degenerate solutions, we impose two explicit constraints during optimization: a lower bound *r*_min_ *>* 0 on the tube radius *r* to prevent the hole from collapsing, and an upper bound *R*_max_ = *σ*_1_(Ƶ) on the major radius, where *σ*_1_ denotes the largest singular value of the centered point cloud, to prevent unrealistically large radii. These bounds are not tuned per dataset; they are fixed relative to the scale of the embedding and applied uniformly across all recordings.

### M.2 Windowed Persistent Homology of Monkey V1 LFP

While torus score and torus error quantify geometric fit quality on a continuous scale, persistent homology provides an independent, coordinate-free verification of the homotopy type of the recovered embedding. We apply a windowed Vietoris–Rips pipeline to assess whether the lag-embedded trajectory recovers the Betti-number signature predicted by the theory (Section 2.2).

#### Preprocessing and embedding

Each channel was preprocessed by 50 Hz notch filtering and 1–50 Hz bandpass filtering, then divided by its Hilbert amplitude envelope (computed in the 1–80 Hz band and low-pass smoothed at 3 Hz) to remove slow amplitude modulation that would otherwise dominate the embedding geometry. The envelope-normalized signal was delay-embedded as

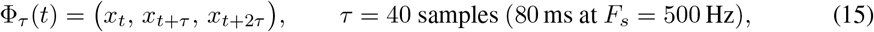

following the delay-embedding framework of Takens [Takens, 2006] and Kantz and Schreiber [Kantz and Schreiber, 2003].

#### Windowed persistence computation

We drew *N* =200 random windows of length 2000 samples (4 s) from the embedded trajectory, sub-sampled each cloud to 300 points to bound the runtime of the two-dimensional persistence computation, and centered and projected each cloud onto the unit sphere so that lifetime thresholds are comparable across windows. For each preprocessed cloud we computed the Vietoris–Rips persistence diagram up to dimension 2 using Ripser [Tralie et al., 2018].

#### Betti-number classification

Each window is summarized by a Betti vector *β* = (*β*_0_, *β*_1_, *β*_2_), where *β*_*k*_ counts the number of dimension-*k* bars whose lifetime (death −birth) exceeds a fixed persistence threshold of 0.30. Since each unit-sphere-projected cloud is connected by construction, *β*_0_ is fixed to 1 for all windows. The window-level distribution over *β* is then aggregated to identify the dominant homotopy type across the recording. The predicted two-torus signature (*β*_0_, *β*_1_, *β*_2_) = (1, 2, 1) is recovered in 46% of windows, with the adjacent configuration (1, 1, 1) in 40% of windows and 14% classified as other, consistent with the theoretical prediction from Section 2.2.

#### Sensitivity analysis

The sensitivity of the window-level Betti distribution to the choice of embedding delay *τ*, window length, sub-sample size, and persistence threshold is reported in Appendix E (Figure 14) and in the sensitivity heatmaps of Figure 4e of the main text.

### M.3 Diagnostic Applied to Real Recordings

Figure 18E shows the Procrustes disparity diagnostic applied to real neural recordings, overlaid on the synthetic calibration distributions from Appendix L.2. Monkey V1 LFP recordings yield a mean Procrustes disparity of 0.1477 (blue vertical line), above the true torus threshold (0.0820, orange dashed line) but well below the bulk of the spurious torus distribution, indicating geometrically valid toroidal structure with moderate noise — consistent with the two dominant oscillatory modes and broadband background observed spectrally in Figure 4. Mouse cortex EEG recordings yield a mean Procrustes disparity of 0.4092 (green vertical line), falling within the spurious torus distribution, reflecting the noisier and less geometrically clean torus structure observed qualitatively in Figure 13. Importantly, this does not indicate failure of the framework: rather, the diagnostic correctly grades geometric quality on a continuous scale, reflecting the degree to which the signal’s oscillatory structure is sustained and spectrally distinct. The mouse EEG result is consistent with the absence of a dominant narrowband peak in its power spectrum (Figure 13c) and with the theoretical prediction that weaker or broader oscillatory modes yield thicker, less geometrically precise tori.

#### Summary

Together, these results establish three points. First, single-channel lag embeddings reflect genuine local oscillatory geometry rather than volume conduction artifacts. Second, the Procrustes disparity diagnostic reliably separates genuine from spurious toroidal geometry on synthetic data. Third, real LFP and EEG recordings fall at interpretable positions on the diagnostic scale, with monkey V1 LFP showing cleaner toroidal geometry than mouse cortex EEG, consistent with their respective spectral structures and the predictions of Section 2.

## N Downstream Decoding: Methods and Additional Results

The decoding analyses use features extracted from fitted lag-embedded point clouds. Each fit supplies 15 features: two major radii and a minor radius describing shape; mean squared error, mean error, and the fraction of points inside the fitted surface describing fit quality; and nine coordinates describing the surface orientation. Those nine coordinates comprise the three-dimensional plane normal and two in-plane axes. The task-stage analysis uses a fitted band half-width in place of the minor radius. These are geometric features, distinct from the Betti numbers used for topological validation.

### N.1 Mouse cortical EEG sleep-state decoding

We divided each recording hour into non-overlapping, two-second windows labeled Wake, NREM, or REM. The EEG was sampled at 400 Hz. For geometric fitting, we removed 60 Hz line noise, filtered the signal from 0.5 to 50 Hz, and normalized its amplitude using a smoothed Hilbert envelope. This normalization was applied before dividing the recording into windows. We then embedded each window in three lag coordinates separated by one sample and fitted an elliptical toroidal surface to the resulting point cloud. Its parameters and orientation supplied the 15 features described above. We evaluated each recording hour separately. Within an hour, we sampled equal numbers of windows from the three states and evaluated a linear discriminant analysis (LDA) classifier using five-fold stratified cross-validation. Twenty-one hours contained enough windows of every state for this analysis. Performance was measured by macro-F1, the mean of the three state-specific F1 scores; we report the mean and standard deviation across hours. Thus, these results summarize separate within-hour decoding analyses across 21 recording hours.

For spectral comparisons, we estimated each window’s power spectrum and computed power in the delta (0.5–4 Hz), theta (4–8 Hz), alpha (8–12 Hz), sigma (12–16 Hz), beta (16–30 Hz), and low-gamma (30–50 Hz) bands. We evaluated each band separately, all six band powers together, and average spectral power over 0.5–50 Hz using the same labels and cross-validation procedure. The geometric features achieved 0.670 ±0.060 macro-F1 across hours, compared with 0.470 ±0.050 for delta power, 0.485± 0.062 for average spectral power, and 0.696 ±0.051 for the combined six-band vector. Table 10 reports the individual-band and geometric decoding scores for each eligible recording hour, showing the variation underlying the aggregate results.

**Table 10:** Mouse EEG sleep-stage decoding by recording hour. Macro-F1 from separate within-hour, five-fold cross-validation analyses. The final row gives mean ± SD across the 21 eligible hours.

| Hour | Delta | Theta | Alpha | Sigma | Beta | Low-γ | Torus |
|---|---|---|---|---|---|---|---|
| 3 | 0.55 | 0.47 | 0.34 | 0.33 | 0.53 | 0.55 | 0.64 |
| 4 | 0.51 | 0.37 | 0.34 | 0.29 | 0.58 | 0.65 | 0.73 |
| 5 | 0.52 | 0.41 | 0.35 | 0.30 | 0.56 | 0.67 | 0.67 |
| 6 | 0.40 | 0.35 | 0.25 | 0.47 | 0.57 | 0.70 | 0.72 |
| 7 | 0.47 | 0.43 | 0.29 | 0.37 | 0.57 | 0.68 | 0.71 |
| 8 | 0.42 | 0.44 | 0.38 | 0.37 | 0.56 | 0.65 | 0.72 |
| 9 | 0.48 | 0.52 | 0.36 | 0.30 | 0.55 | 0.74 | 0.74 |
| 10 | 0.51 | 0.45 | 0.37 | 0.31 | 0.54 | 0.68 | 0.69 |
| 11 | 0.44 | 0.49 | 0.37 | 0.33 | 0.55 | 0.60 | 0.67 |
| 12 | 0.41 | 0.54 | 0.29 | 0.33 | 0.45 | 0.59 | 0.60 |
| 13 | 0.47 | 0.39 | 0.34 | 0.33 | 0.54 | 0.59 | 0.64 |
| 15 | 0.48 | 0.46 | 0.34 | 0.49 | 0.67 | 0.68 | 0.74 |
| 16 | 0.44 | 0.40 | 0.28 | 0.35 | 0.56 | 0.68 | 0.75 |
| 17 | 0.43 | 0.32 | 0.32 | 0.34 | 0.57 | 0.65 | 0.68 |
| 18 | 0.42 | 0.38 | 0.39 | 0.33 | 0.61 | 0.68 | 0.72 |
| 19 | 0.41 | 0.35 | 0.33 | 0.35 | 0.46 | 0.54 | 0.57 |
| 20 | 0.58 | 0.39 | 0.41 | 0.35 | 0.55 | 0.64 | 0.58 |
| 21 | 0.51 | 0.44 | 0.36 | 0.29 | 0.54 | 0.64 | 0.68 |
| 22 | 0.45 | 0.35 | 0.31 | 0.30 | 0.47 | 0.56 | 0.57 |
| 23 | 0.45 | 0.35 | 0.33 | 0.32 | 0.55 | 0.66 | 0.61 |
| 24 | 0.53 | 0.46 | 0.32 | 0.34 | 0.54 | 0.56 | 0.64 |
| Mean ± SD | 0.47 ± 0.05 | 0.42 ± 0.06 | 0.34 ± 0.04 | 0.34 ± 0.05 | 0.55 ± 0.05 | 0.64 ± 0.05 | 0.67 ± 0.06 |

To asses whether decoding remains effective when AR order is misestimated, we compare the full 15-feature decoder with an 11-feature version retaining only the two major radii and nine orientation coordinates. The reduced feature set achieved 0.651± 0.072 macro-F1, compared with 0.670 ± 0.060 for the full feature set. The paired difference across hours was not significant (*p* = 0.0822). Thus, although order-recovery errors can affect the inferred torus dimension and discrete Betti classification, we do not observe a statistically reliable degradation in downstream decoding performance.

### N.2 Macaque reach-direction decoding

We analyzed motor-cortex LFP from two macaques performing the delayed reaching task. For each trial, we extracted a one-second segment from 250 ms before to 750 ms after movement onset. Each segment was linearly detrended, normalized by its median and median absolute deviation, filtered from 2 to 55 Hz, and delay-embedded. We selected the embedding delay and dimension once for each LFP recording from its movement-aligned signal. Embeddings with more than three coordinates were projected onto their leading three principal coordinates. We fitted an elliptical toroidal surface to at most 300 sampled points from each trial’s cloud and extracted 15 geometric features for reach-direction decoding.

We decoded the six reach directions separately within each LFP recording. Trials were balanced across directions, and an LDA classifier was evaluated using five-fold stratified cross-validation. Spectral comparisons used individual band powers, average spectral power, and the combined band-power vector. In total, 237 LFP recordings contained all six directions: 135 from Monkey M and 102 from Monkey T. Across these recordings, geometric features achieved 0.181± 0.032 macro-F1, compared with 0.157 ± 0.030 for beta-band power and 0.159 ± 0.030 for average spectral power. The combined band-power vector achieved approximately 0.212 macro-F1. Table 11 reports the results for each macaque separately. Each result comes from training and evaluating within a recording.

**Table 11:** Reach-direction decoding by macaque. Mean ±SD macro-F1 across LFP recordings with all six directions. Models were trained and evaluated separately within each recording.

| Macaque | LFPs | Torus features | Beta power | Avg. PSD |
|---|---|---|---|---|
| Monkey M | 135 | 0.175 ± 0.034 | 0.155 ± 0.030 | 0.151 ± 0.027 |
| Monkey T | 102 | 0.188 ± 0.028 | 0.159 ± 0.026 | 0.169 ± 0.031 |

### N.3 Macaque task-stage decoding

We used one Monkey T recording to test whether the geometric features also distinguish events within a reaching trial. The analysis included 198 short-delay trials, with 33 trials for each reach direction. From every trial we extracted six non-overlapping, 300-ms windows: immediately before and after the temporal cue, immediately before and after the spatial cue, and immediately before and after the go signal. Post-cue windows began after the corresponding cue ended. This yielded 1,188 labeled windows.

Each window was detrended, normalized by its median and median absolute deviation, and filtered from 2 to 55 Hz. A three-coordinate delay embedding with a 3-ms delay yielded 294 points per window. We fitted the resulting point cloud and extracted the 15 geometric features described above; 1,177 of the 1,188 fits were usable. For five-fold LDA, all six windows from a given trial were assigned to the same fold. Missing fit features were filled using medians calculated from the training fold, and feature scaling was learned from that fold.

Geometry alone achieved 0.293 macro-F1. Beta-band power alone achieved 0.291, rising to 0.347 when combined with the geometric features; average spectral power rose from 0.194 to 0.311 when combined with them. A bootstrap resampling whole trials gave a 95% interval of 0.268–0.318 for geometry-only macro-F1. A within-trial permutation test of stage labels gave *p* = 0.0010. This is an exploratory, within-recording analysis of task-stage information.

### N.4 Mouse hippocampal LFP processing and geometric features

We analyzed hippocampal LFP recordings from one mouse using the supplied channel-quality annotations and behavioral labels. The decoding comparison used windows labeled immobile or sleep; mobile and unlabeled periods were excluded. The raw signals were sampled at 20 kHz. We extracted two-second windows at one-second intervals, removed 60 Hz line noise, filtered each window from 1 to 80 Hz, and resampled it to 500 Hz. We normalized amplitude using a Hilbert envelope smoothed with a 3 Hz low-pass filter, then scaled each window to [− 1, 1] for geometric fitting.

We selected a delay separately for each channel recording from the autocorrelation of sampled two-second windows. For each sampled window, we selected the first non-positive autocorrelation value within the delay search range, or the value closest to zero when no crossing occurred. The median selected delay was then held fixed across windows from that recording. Each window was embedded in three lag coordinates and fitted with an elliptical toroidal surface. The fit supplied the 15 geometric features described at the beginning of this appendix. Behavioral labels were assigned at the window center.

### N.5 Mouse hippocampal LFP decoding and spectral comparisons

We compared the geometric features with theta-band power and average spectral power. Spectra were estimated from the processed windows using Welch’s method with a Hann window. Theta power was integrated over 4–8 Hz and log-transformed. The average-PSD feature was the mean log spectral density at 1 Hz intervals over 1–80 Hz. Each spectral comparison therefore used one feature, whereas the geometric decoder used all 15 fitted features.

We performed 100 repeated within-recording decoding analyses. In each repeat, we randomly selected an eligible channel recording from the mouse’s recording days and sampled 50 windows per state, using replacement when fewer windows were available. Within each state, 32 sampled windows were assigned to training and 18 to testing, giving 64 training and 36 test windows. All feature sets used the same sampled windows and splits. We fitted an *L*_2_-regularized logistic regression classifier, with median imputation and feature standardization learned from the training set. Performance was measured by macro-F1, the mean of the immobile and sleep F1 scores.

Across the 100 repeats, geometric features achieved 0.624± 0.099 macro-F1, compared with 0.592± 0.124 for theta power and 0.532± 0.111 for average PSD (mean± SD). Two-sided paired Wilcoxon signed-rank tests across matched repeats gave *p* = 0.0278 versus theta power and *p* = 3.52 × 10^*−*7^ versus average PSD, corresponding to the mouse hippocampal LFP row in Table 4.

## NeurIPS Paper Checklist

The checklist is designed to encourage best practices for responsible machine learning research, addressing issues of reproducibility, transparency, research ethics, and societal impact. Do not remove the checklist: **The papers not including the checklist will be desk rejected**. The checklist should follow the references and follow the (optional) supplemental material. The checklist does NOT count towards the page limit.

Please read the checklist guidelines carefully for information on how to answer these questions. For each question in the checklist:

- You should answer [Yes], [No], or [N/A].
- [N/A] means either that the question is Not Applicable for that particular paper or the relevant information is Not Available.
- Please provide a short (1–2 sentence) justification right after your answer (even for [N/A]).

### The checklist answers are an integral part of your paper submission

They are visible to the reviewers, area chairs, senior area chairs, and ethics reviewers. You will also be asked to include it (after eventual revisions) with the final version of your paper, and its final version will be published with the paper.

The reviewers of your paper will be asked to use the checklist as one of the factors in their evaluation. While [Yes] is generally preferable to [No], it is perfectly acceptable to answer [No] provided a proper justification is given (e.g., error bars are not reported because it would be too computationally expensive” or “we were unable to find the license for the dataset we used”). In general, answering [No] or [N/A] is not grounds for rejection. While the questions are phrased in a binary way, we acknowledge that the true answer is often more nuanced, so please just use your best judgment and write a justification to elaborate. All supporting evidence can appear either in the main paper or the supplemental material, provided in appendix. If you answer [Yes] to a question, in the justification please point to the section(s) where related material for the question can be found.

IMPORTANT, please:

- **Delete this instruction block, but keep the section heading “NeurIPS Paper Checklist”**,
- **Keep the checklist subsection headings, questions/answers and guidelines below**.
- **Do not modify the questions and only use the provided macros for your answers**.

#### 1. Claims

Question: Do the main claims made in the abstract and introduction accurately reflect the paper’s contributions and scope?

Answer: [Yes]

Justification: The abstract and Section 1 identify the central contributions: an idealized *K*-torus characterization of oscillatory lag embeddings, a local TVAR approximation, DeepLagField, and empirical geometry and decoding tests. Sections 2–4 state the principal theoretical conditions and report the mixed window-level topological findings.

Guidelines:

- The answer [N/A] means that the abstract and introduction do not include the claims made in the paper.
- The abstract and/or introduction should clearly state the claims made, including the contributions made in the paper and important assumptions and limitations. A [No] or [N/A] answer to this question will not be perceived well by the reviewers.
- The claims made should match theoretical and experimental results, and reflect how much the results can be expected to generalize to other settings.
- It is fine to include aspirational goals as motivation as long as it is clear that these goals are not attained by the paper.

#### 2. Limitations

Question: Does the paper discuss the limitations of the work performed by the authors?

Answer: [Yes]

Justification: The introduction and Section 2.2 distinguish the idealized noiseless, nonresonant setting from noisy and damped recordings. Section 3.5 reports incomplete toroidal recovery across windows and poorer geometric fits in mouse EEG. Appendix C.1 discusses the window-length tradeoff, Appendix E gives representative limitation cases, and Appendix K.2 restricts scalability claims to the tested network sizes.

Guidelines:

- The answer [N/A] means that the paper has no limitation while the answer [No] means that the paper has limitations, but those are not discussed in the paper.
- The authors are encouraged to create a separate “Limitations” section in their paper.
- The paper should point out any strong assumptions and how robust the results are to violations of these assumptions (e.g., independence assumptions, noiseless settings, model well-specification, asymptotic approximations only holding locally). The authors should reflect on how these assumptions might be violated in practice and what the implications would be.
- The authors should reflect on the scope of the claims made, e.g., if the approach was only tested on a few datasets or with a few runs. In general, empirical results often depend on implicit assumptions, which should be articulated.
- The authors should reflect on the factors that influence the performance of the approach. For example, a facial recognition algorithm may perform poorly when image resolution is low or images are taken in low lighting. Or a speech-to-text system might not be used reliably to provide closed captions for online lectures because it fails to handle technical jargon.
- The authors should discuss the computational efficiency of the proposed algorithms and how they scale with dataset size.
- If applicable, the authors should discuss possible limitations of their approach to address problems of privacy and fairness.
- While the authors might fear that complete honesty about limitations might be used by reviewers as grounds for rejection, a worse outcome might be that reviewers discover limitations that aren’t acknowledged in the paper. The authors should use their best judgment and recognize that individual actions in favor of transparency play an important role in developing norms that preserve the integrity of the community. Reviewers will be specifically instructed to not penalize honesty concerning limitations.

#### 3. Theory assumptions and proofs

Question: For each theoretical result, does the paper provide the full set of assumptions and a complete (and correct) proof?

Answer: [Yes]

Justification: The assumptions and proofs supporting the theoretical results are provided in Appendices B and C. In particular, Lemmas B.1–B.4 and Proposition B.5 establish the connection between oscillatory AR modes and the K-torus geometry under the stated idealized assumptions. Appendix C derives the local approximation bound for time-varying AR processes and discusses the continuity of polynomial roots under coefficient perturbations. The main text summarizes the theoretical results and refers to their detailed derivations.

Guidelines:

- The answer [N/A] means that the paper does not include theoretical results.
- All the theorems, formulas, and proofs in the paper should be numbered and cross-referenced.
- All assumptions should be clearly stated or referenced in the statement of any theorems.
- The proofs can either appear in the main paper or the supplemental material, but if they appear in the supplemental material, the authors are encouraged to provide a short proof sketch to provide intuition.
- Inversely, any informal proof provided in the core of the paper should be complemented by formal proofs provided in appendix or supplemental material.
- Theorems and Lemmas that the proof relies upon should be properly referenced.

#### 4. Experimental result reproducibility

Question: Does the paper fully disclose all the information needed to reproduce the main experimental results of the paper to the extent that it affects the main claims and/or conclusions of the paper (regardless of whether the code and data are provided or not)?

Answer: [Yes]

Justification: The experimental methodology and evaluation procedures are documented in Sections 3–4 and Appendices G, I, K, M, and N. These include the model architecture, physics-informed training framework, data preprocessing, synthetic benchmarks, computational setup, topological validation, and downstream decoding protocols. Implementation details, including the specific regularization choices and hyperparameters, are provided in the publicly available code repository.

Guidelines:

- The answer [N/A] means that the paper does not include experiments.
- If the paper includes experiments, a [No] answer to this question will not be perceived well by the reviewers: Making the paper reproducible is important, regardless of whether the code and data are provided or not.
- If the contribution is a dataset and/or model, the authors should describe the steps taken to make their results reproducible or verifiable.
- Depending on the contribution, reproducibility can be accomplished in various ways. For example, if the contribution is a novel architecture, describing the architecture fully might suffice, or if the contribution is a specific model and empirical evaluation, it may be necessary to either make it possible for others to replicate the model with the same dataset, or provide access to the model. In general. releasing code and data is often one good way to accomplish this, but reproducibility can also be provided via detailed instructions for how to replicate the results, access to a hosted model (e.g., in the case of a large language model), releasing of a model checkpoint, or other means that are appropriate to the research performed.
- While NeurIPS does not require releasing code, the conference does require all submissions to provide some reasonable avenue for reproducibility, which may depend on the nature of the contribution. For example
  a. If the contribution is primarily a new algorithm, the paper should make it clear how to reproduce that algorithm.
  b. If the contribution is primarily a new model architecture, the paper should describe the architecture clearly and fully.
  c. If the contribution is a new model (e.g., a large language model), then there should either be a way to access this model for reproducing the results or a way to reproduce the model (e.g., with an open-source dataset or instructions for how to construct the dataset).
  d. We recognize that reproducibility may be tricky in some cases, in which case authors are welcome to describe the particular way they provide for reproducibility. In the case of closed-source models, it may be that access to the model is limited in some way (e.g., to registered users), but it should be possible for other researchers to have some path to reproducing or verifying the results.

#### 5. Open access to data and code

Question: Does the paper provide open access to the data and code, with sufficient instructions to faithfully reproduce the main experimental results, as described in supplemental material?

Answer: [No]

Justification: The complete implementation, documentation, and tutorials are publicly available at https://github.com/erfanzabeh/NeuralFieldManifold under the MIT license. Some of the experimental datasets are also publicly available and their original sources are appropriately referenced in the manuscript. However, certain electrophysiological recordings are not publicly available for redistribution.

Guidelines:

- The answer [N/A] means that paper does not include experiments requiring code.
- Please see the NeurIPS code and data submission guidelines (https://neurips.cc/public/guides/CodeSubmissionPolicy) for more details.
- While we encourage the release of code and data, we understand that this might not be possible, so [No] is an acceptable answer. Papers cannot be rejected simply for not including code, unless this is central to the contribution (e.g., for a new open-source benchmark).
- The instructions should contain the exact command and environment needed to run to reproduce the results. See the NeurIPS code and data submission guidelines (https://neurips.cc/public/guides/CodeSubmissionPolicy) for more details.
- The authors should provide instructions on data access and preparation, including how to access the raw data, preprocessed data, intermediate data, and generated data, etc.
- The authors should provide scripts to reproduce all experimental results for the new proposed method and baselines. If only a subset of experiments are reproducible, they should state which ones are omitted from the script and why.
- At submission time, to preserve anonymity, the authors should release anonymized versions (if applicable).
- Providing as much information as possible in supplemental material (appended to the paper) is recommended, but including URLs to data and code is permitted.

#### 6. Experimental setting/details

Question: Does the paper specify all the training and test details (e.g., data splits, hyperparameters, how they were chosen, type of optimizer) necessary to understand the results?

Answer: [Yes]

Justification: The experimental settings are detailed in Appendices G, I, K, M, and N, including dataset preparation, model architecture, training procedures, optimization settings, computational resources, and evaluation protocols. The training and testing procedures, including data splits, cross-validation, and statistical analyses, are described for the different experiments. Additional implementation details and hyperparameters are available in the publicly released code repository.

Guidelines:

- The answer [N/A] means that the paper does not include experiments.
- The experimental setting should be presented in the core of the paper to a level of detail that is necessary to appreciate the results and make sense of them.
- The full details can be provided either with the code, in appendix, or as supplemental material.

#### 7. Experiment statistical significance

Question: Does the paper report error bars suitably and correctly defined or other appropriate information about the statistical significance of the experiments?

Answer: [Yes]

±

±

Justification: The main results report variability and significance using specified units: Table 3 gives mean ± SEM across window-level block bootstraps; Table 4 gives mean ± SD of decoding F1 scores and paired test results. Figure 6 and Appendix N provide cross-validation procedures, bootstrap confidence intervals, permutation tests, and paired comparisons. Appendix N distinguishes the 21 recording-hour, 237 recording, and 100 repeated-split summaries. Guidelines:

- The answer [N/A] means that the paper does not include experiments.
- The authors should answer [Yes] if the results are accompanied by error bars, confidence intervals, or statistical significance tests, at least for the experiments that support the main claims of the paper.
- The factors of variability that the error bars are capturing should be clearly stated (for example, train/test split, initialization, random drawing of some parameter, or overall run with given experimental conditions).
- The method for calculating the error bars should be explained (closed form formula, call to a library function, bootstrap, etc.)
- The assumptions made should be given (e.g., Normally distributed errors).
- It should be clear whether the error bar is the standard deviation or the standard error of the mean.
- It is OK to report 1-sigma error bars, but one should state it. The authors should preferably report a 2-sigma error bar than state that they have a 96% CI, if the hypothesis of Normality of errors is not verified.
- For asymmetric distributions, the authors should be careful not to show in tables or figures symmetric error bars that would yield results that are out of range (e.g., negative error rates).
- If error bars are reported in tables or plots, the authors should explain in the text how they were calculated and reference the corresponding figures or tables in the text.

#### 8. Experiments compute resources

Question: For each experiment, does the paper provide sufficient information on the computer resources (type of compute workers, memory, time of execution) needed to reproduce the experiments?

Answer: [Yes]

Justification: Appendix K.1 reports the workstation configuration (RTX 5090 GPUs, Threadripper CPU, 512 GB RAM, and Ubuntu), default model size, optimizer, batch size, epochs, and per-step timing. Appendix K.2 gives model-width scaling and inference timing.

Guidelines:

- The answer [N/A] means that the paper does not include experiments.
- The paper should indicate the type of compute workers CPU or GPU, internal cluster, or cloud provider, including relevant memory and storage.
- The paper should provide the amount of compute required for each of the individual experimental runs as well as estimate the total compute.
- The paper should disclose whether the full research project required more compute than the experiments reported in the paper (e.g., preliminary or failed experiments that didn’t make it into the paper).

#### 9. Code of ethics

Question: Does the research conducted in the paper conform, in every respect, with the NeurIPS Code of Ethics https://neurips.cc/public/EthicsGuidelines?

Answer: [Yes]

Justification: Ethical considerations have been taken into account and the research conforms fully to the NeurIPS code of ethics.

Guidelines:

- The answer [N/A] means that the authors have not reviewed the NeurIPS Code of Ethics.
- If the authors answer [No], they should explain the special circumstances that require a deviation from the Code of Ethics.
- The authors should make sure to preserve anonymity (e.g., if there is a special consideration due to laws or regulations in their jurisdiction).

#### 10. Broader impacts

Question: Does the paper discuss both potential positive societal impacts and negative societal impacts of the work performed?

Answer: [No]

Justification: The introduction and discussion motivate potential neuroscience, clinical, and brain–computer interface benefits, but the manuscript does not explicitly discuss both positive and potential negative societal impacts.

Guidelines:

- The answer [N/A] means that there is no societal impact of the work performed.
- If the authors answer [N/A] or [No], they should explain why their work has no societal impact or why the paper does not address societal impact.
- Examples of negative societal impacts include potential malicious or unintended uses (e.g., disinformation, generating fake profiles, surveillance), fairness considerations (e.g., deployment of technologies that could make decisions that unfairly impact specific groups), privacy considerations, and security considerations.
- The conference expects that many papers will be foundational research and not tied to particular applications, let alone deployments. However, if there is a direct path to any negative applications, the authors should point it out. For example, it is legitimate to point out that an improvement in the quality of generative models could be used to generate Deepfakes for disinformation. On the other hand, it is not needed to point out that a generic algorithm for optimizing neural networks could enable people to train models that generate Deepfakes faster.
- The authors should consider possible harms that could arise when the technology is being used as intended and functioning correctly, harms that could arise when the technology is being used as intended but gives incorrect results, and harms following from (intentional or unintentional) misuse of the technology.
- If there are negative societal impacts, the authors could also discuss possible mitigation strategies (e.g., gated release of models, providing defenses in addition to attacks, mechanisms for monitoring misuse, mechanisms to monitor how a system learns from feedback over time, improving the efficiency and accessibility of ML).

#### 11. Safeguards

Question: Does the paper describe safeguards that have been put in place for responsible release of data or models that have a high risk for misuse (e.g., pre-trained language models, image generators, or scraped datasets)?

Answer: [N/A]

Justification: This paper poses no such risks.

Guidelines:

- The answer [N/A] means that the paper poses no such risks.
- Released models that have a high risk for misuse or dual-use should be released with necessary safeguards to allow for controlled use of the model, for example by requiring that users adhere to usage guidelines or restrictions to access the model or implementing safety filters.
- Datasets that have been scraped from the Internet could pose safety risks. The authors should describe how they avoided releasing unsafe images.
- We recognize that providing effective safeguards is challenging, and many papers do not require this, but we encourage authors to take this into account and make a best faith effort.

#### 12. Licenses for existing assets

Question: Are the creators or original owners of assets (e.g., code, data, models), used in the paper, properly credited and are the license and terms of use explicitly mentioned and properly respected?

Answer: [Yes]

Justification: Appendix G credits the original studies and describes the biological recording sources; the references also credit methodological software.

Guidelines:

- The answer [N/A] means that the paper does not use existing assets.
- The authors should cite the original paper that produced the code package or dataset.
- The authors should state which version of the asset is used and, if possible, include a URL.
- The name of the license (e.g., CC-BY 4.0) should be included for each asset.
- For scraped data from a particular source (e.g., website), the copyright and terms of service of that source should be provided.
- If assets are released, the license, copyright information, and terms of use in the package should be provided. For popular datasets, paperswithcode.com/datasets has curated licenses for some datasets. Their licensing guide can help determine the license of a dataset.
- For existing datasets that are re-packaged, both the original license and the license of the derived asset (if it has changed) should be provided.
- If this information is not available online, the authors are encouraged to reach out to the asset’s creators.

#### 13. New assets

Question: Are new assets introduced in the paper well documented and is the documentation provided alongside the assets?

Answer: [Yes]

Justification: The main new asset is the publicly released NeuralFieldMani-fold/DeepLagField software package. The manuscript links the code and project tutorials, and the public repository provides installation guidance, API documentation, examples, and an MIT license. No new independent biological dataset is claimed as a released contribution.

Guidelines:

- The answer [N/A] means that the paper does not release new assets.
- Researchers should communicate the details of the dataset/code/model as part of their submissions via structured templates. This includes details about training, license, limitations, etc.
- The paper should discuss whether and how consent was obtained from people whose asset is used.
- At submission time, remember to anonymize your assets (if applicable). You can either create an anonymized URL or include an anonymized zip file.

#### 14. Crowdsourcing and research with human subjects

Question: For crowdsourcing experiments and research with human subjects, does the paper include the full text of instructions given to participants and screenshots, if applicable, as well as details about compensation (if any)?

Answer: [N/A]

Justification: This paper does not involve such experiments.

Guidelines:

- The answer [N/A] means that the paper does not involve crowdsourcing nor research with human subjects.
- Including this information in the supplemental material is fine, but if the main contribution of the paper involves human subjects, then as much detail as possible should be included in the main paper.
- According to the NeurIPS Code of Ethics, workers involved in data collection, curation, or other labor should be paid at least the minimum wage in the country of the data collector.

#### 15. Institutional review board (IRB) approvals or equivalent for research with human subjects

Question: Does the paper describe potential risks incurred by study participants, whether such risks were disclosed to the subjects, and whether Institutional Review Board (IRB) approvals (or an equivalent approval/review based on the requirements of your country or institution) were obtained?

Answer: [N/A]

Justification: This paper does not involve any user studies.

Guidelines:

- The answer [N/A] means that the paper does not involve crowdsourcing nor research with human subjects.
- Depending on the country in which research is conducted, IRB approval (or equivalent) may be required for any human subjects research. If you obtained IRB approval, you should clearly state this in the paper.
- We recognize that the procedures for this may vary significantly between institutions and locations, and we expect authors to adhere to the NeurIPS Code of Ethics and the guidelines for their institution.
- For initial submissions, do not include any information that would break anonymity (if applicable), such as the institution conducting the review.

#### 16. Declaration of LLM usage

Question: Does the paper describe the usage of LLMs if it is an important, original, or non-standard component of the core methods in this research? Note that if the LLM is used only for writing, editing, or formatting purposes and does *not* impact the core methodology, scientific rigor, or originality of the research, declaration is not required.

Answer: [N/A]

Justification: No large language model is presented as an important, original, or non-standard component of the scientific method. The proposed estimators, mathematical analysis, and decoding experiments do not require LLMs.

Guidelines:

- The answer [N/A] means that the core method development in this research does not involve LLMs as any important, original, or non-standard components.
- Please refer to our LLM policy in the NeurIPS handbook for what should or should not be described.

## Notes

https://zabehlab.github.io/NeuralFieldManifold/

https://github.com/ZabehLab/NeuralFieldManifold

### Competing Interest Statement

The authors have declared no competing interest.

